# Astrocyte-like subpopulation of NG2 glia in the adult mouse cortex exhibits characteristics of neural progenitor cells and is capable of forming neuron-like cells after ischemic injury

**DOI:** 10.1101/2023.02.20.529180

**Authors:** Lucie Janeckova, Tomas Knotek, Jan Kriska, Zuzana Hermanova, Denisa Kirdajova, Jan Kubovciak, Linda Berkova, Jana Tureckova, Sara Camacho Garcia, Katerina Galuskova, Michal Kolar, Miroslava Anderova, Vladimir Korinek

## Abstract

Glia cells expressing neuron-glial antigen 2 (NG2) play a critical role as oligodendrocyte precursor cells (OPCs) in the healthy brain; however, their differentiation potential after ischemic injury remains an unresolved question. Here, we aimed to elucidate the heterogeneity and role of NG2 glia in the ischemic brain. We used transgenic mice to label NG2-expressing cells and their progeny with red fluorescent protein tdTomato in the healthy brains and those after focal cerebral ischemia (FCI). Based on single-cell RNA sequencing, the labeled glial cells were divided into five distinct subpopulations. The identity of these subpopulations was determined based on gene expression patterns. In addition, membrane properties were further analyzed using the patch-clamp technique. Three of the observed subpopulations represented OPCs, whereas the fourth group exhibited characteristics of cells destined for oligodendrocyte fate. The fifth subpopulation of NG2 glia carried astrocytic markers. Importantly, we detected features of neural progenitors in these cells. This subpopulation was present in both healthy and post-ischemic tissue; however, its gene expression changed after ischemia, with genes related to neurogenesis being more abundant. Neurogenic gene expression was monitored over time and complemented by immunohistochemical staining, which showed increased numbers of Purkinje cell protein 4-positive NG2 cells at the edge of the ischemic lesion 12 days after FCI, and NeuN-positive NG2 cells 28 days after injury, indicating the existence of neuron-like cells that develop from NG2 glia in the ischemic tissue. Our results provide further insight into the differentiation plasticity and neurogenic potential of NG2 glia after stroke.

**Main Points:**

- Different subpopulations of NG2 glia in the healthy and ischemic adult cortex were identified based on their gene expression and membrane properties.
- Astrocyte-like NG2 glia exhibit neurogenic gene expression and are more abundant in post-ischemic tissue.
- Progeny of NG2-positive cells carrying neuronal marker NeuN was observed at the edge of the ischemic lesion.

## INTRODUCTION

NG2 glia or polydendrocytes are central nervous system (CNS) glial cells that can be recognized by the presence of chondroitin sulfate proteoglycan 4 (Cspg4, also called neuron-glial antigen 2, NG2) and platelet-derived growth factor receptor alpha (Pdgfrα) (reviewed in (Nishiyama et al. 2016)). In the adult cerebral cortex, NG2 glia account for about 70% of proliferating cells. Thus, they are the most proliferating cell type outside the neurogenic zones: the subventricular zone and the dentate gyrus (Dimou et al. 2008). NG2 glia are present throughout the CNS parenchyma and tend to be evenly distributed (Nishiyama, Chang, and Trapp 1999). NG2 glia, also known as oligodendrocyte precursor cells (OPCs), are known for their ability to form mature oligodendrocytes both during development and in the adult brain (Young et al. 2013, Zhu et al. 2011) not only under physiological but also under pathological conditions (Wilson, Scolding, and Raine 2006). In addition, direct synaptic communication between NG2 glia and neurons has been described: These cells receive both excitatory and inhibitory inputs (Bergles et al. 2000, Lin and Bergles 2004). Moreover, glutamatergic synapses between neurons and NG2 glia can undergo long-term potentiation, suggesting an important role in the membrane properties of this type of glia (Ge et al. 2006). In particular, glutamatergic signaling plays an essential role in regulating the migration of NG2 glia and their differentiation into oligodendrocytes (Li et al. 2020). In addition to their role as progenitor cells, NG2 glia also perform other functions: they contribute to maintaining the integrity of the blood-brain barrier, have immunomodulatory properties, and play a role in various CNS diseases (reviewed in (Akay, Effenberger, and Tsai 2021)).

Interestingly, the differentiation potential of NG2 glia may not be limited to oligodendrogenesis. Early *in vitro* studies have shown that the formation of astrocytes from NG2 glia can be induced by various morphogens and growth factors (Lillien, Sendtner, and Raff 1990, Raff, Miller, and Noble 1983) such as interferon gamma (IFNγ) (Tanner, Cherry, and Mayer-Proschel 2011). However, the results of *in vivo* fate-mapping studies suggest that astrogliogenesis from NG2 glia in the healthy brain is restricted to prenatal development (Huang et al. 2014, Zhu et al. 2011, Zhu, Bergles, and Nishiyama 2008). In addition, several studies have revealed an intriguing but controversial possibility of neurogenesis in different brain regions (Dayer et al. 2005, Guo et al. 2010, Rivers et al. 2008, Tamura et al. 2007). However, the results of other fate-mapping studies did not confirm this phenomenon (Clarke et al. 2012, Dimou et al. 2008, Kang et al. 2010). Thus, the differentiation potential of these cells in the adult brain appears to be limited to oligodendrogenesis under physiological conditions.

Importantly, NG2 glia are involved in tissue regeneration in various brain injuries, including ischemic injury (Hernandez et al. 2021, Song et al. 2017). Ischemic stroke is the leading cause of disability and death worldwide (Katan and Luft 2018, Virani et al. 2021) and treatment options for this disease are still very limited (reviewed in (Rabinstein 2017)). After the ischemic injury, NG2-expressing cells increase in number in the ischemic penumbra, their morphology changes, and together with astrocytes, they contribute to glial scar formation (reviewed in (Adams and Gallo 2018)). A number of previous studies have demonstrated the heterogeneity of NG2 glia and its functions in the recovery process. For example, after an ischemic stroke, an increase in the density of the NG2 glia subpopulation expressing GPR17 was observed (Bonfanti et al. 2017). This subpopulation acts as a reservoir of OPCs and is activated after injury (Boda et al. 2011, Bonfanti et al. 2017, Vigano et al. 2016). In addition, a subset of cells in the NG2 lineage has been found to express astrocyte markers to varying degrees after different types of injury (Hackett et al. 2018). NG2 glia expressing glial fibrillary acidic protein (GFAP), a typical marker for reactive astrocytes, have recently been described in the post-ischemic cortex (Kirdajova et al. 2021, Valny et al. 2018). A small number of studies have also described the differentiation of NG2 glia into neural progenitor cells after brain injury, based on their electrophysiological properties and the presence of neuroblast marker doublecortin (Dcx) (Honsa et al. 2012, Zawadzka et al. 2010). However, in other studies, no colocalization of neuronal markers with NG2 glia was observed after injury (discussed in (Kirdajova and Anderova 2020)). Thus, the exact role and differentiation potential of NG2 glia after adult brain injury remains unclear, and further research is needed. Nonetheless, the study of NG2 glia and their role in ischemia may contribute to a much-needed understanding of ischemic stroke and the mechanisms of subsequent regeneration.

In this study, we investigated the role of NG2 glia in the adult mouse brain after focal cerebral ischemia (FCI). We used single-cell RNA sequencing (scRNA-seq) in combination with genetic labeling and cell tracking to characterize the NG2 cell line and its progeny in detail. In addition, gene expression analysis was combined with immunohistochemical analysis of the cerebral cortex at different time points after injury. The patch-clamp technique was used to functionally validate different cell subtypes defined by gene expression profiles. Five subpopulations of NG2 glia were distinguished that were also present in the healthy cortex and whose presence, distribution, and expression of several marker genes changed after FCI. In addition to cells representing the OPC lineage, we identified a subpopulation previously described as “astrocyte-like NG2 glia” with similarities to neural progenitors. In contrast to previous studies, we also detected this cell type in healthy tissue, but its gene expression was markedly different from that in the ischemic cortex, mainly due to the absence of GFAP expression in NG2 glia of the healthy brain. Interestingly, several genes related to neurogenesis were expressed during cortical regeneration after FCI. Finally, sparse NG2 glia with a marker of mature neurons were detected in the injured area 28 days after ischemia.

## MATERIAL AND METHODS

### Transgenic animals

All procedures involving the use of laboratory mice were performed in accordance with the European Communities Council Directive of November 24, 1986 (86/609/EEC) and the Animal Welfare Guidelines approved by the Institute of Experimental Medicine of the Czech Academy of Sciences (approval numbers 18/2011, 146/2013, 91/2016, 62/2017, 2/2017, and 10-2021-P). Mouse strains B6;129S6-Gt(ROSA)26Sor^tm14(CAG-tdTomato)Hze^/J and B6.Cg-Tg(Cspg4-cre/Esr1*)BAkik/J were purchased from the Jackson Laboratory (JAX stock #007908 and #008538, respectively). Young adult male mice (P59 - P69 at the beginning of the experiment) were used in this study. DNA recombination was achieved by two intraperitoneal (i.p.) injections of tamoxifen (200 mg/kg animal body weight; purchased from Toronto Research Chemicals, Toronto, Ontario, Canada) dissolved in corn oil (Sigma-Aldrich, St. Louis, MO, USA) on two consecutive days. The following treatments (listed below) were administered on day 15 after the second tamoxifen injection.

### Induction of focal cerebral ischemia (FCI)

A well-established model of permanent middle cerebral artery occlusion (MCAO) was used to induce FCI (Colak, Filiano, and Johnson 2011). The entire procedure has been described in detail previously (Kriska et al. 2021). A group of mice that underwent this procedure was designated ‘MCAO’. In a control group (sham-operated animals; referred to as ‘CTRL’), the same procedure was used, but the vessel was not occluded. This distal model of MCAO has a high survival rate (> 95%) and good reproducibility because it typically results in an infarct lesion of relatively small volume mostly in the cortical region (Androvic et al. 2020).

### Tissue isolation and preparation of single-cell suspension

All animals were deeply anesthetized with pentobarbital solution (PTB; 100 mg/kg, i.p.; Sigma-Aldrich) prior to tissue isolation. The preparation of tissue sections for immunohistochemical analysis was described earlier (Kriska et al. 2021). To prepare coronal brain sections for patch-clamp recordings, anesthetized animals were perfused transcardially with ice-cold isolation NMDG buffer containing (in mM): 110 N-methyl-D-glucamine (NMDG)-Cl, 2.5 KCl, 24.5 NaHCO_3_, 1.25 Na_2_HPO_4_, 0.5 CaCl_2_, 7 MgCl_2_, 20 glucose, osmolality 290 ± 3 mOsmol/kg. The isolation buffer was gassed with 5 % CO_2_ to maintain pH 7.4. Subsequently, the experimental animals were decapitated, the brains were quickly removed from the skull and cut into 200 µm thick coronal slices using a Leica vibratome VT 1200S (Leica Biosystems, Buffalo, IL, USA). The slices were then incubated for 30 minutes at 34 °C in the isolation NMDG buffer and for another 30 minutes in artificial cerebrospinal fluid (aCSF) containing (in mM): 122 NaCl, 3 KCl, 1.5 CaCl_2_, 1.3 MgCl_2_, 1.25 Na_2_HPO_4_, 28 NaHCO_3_, and 10 D-glucose (osmolality 300 ± 5 mOsmol/kg). Both incubation solutions were gassed with 5 % CO_2_.

To obtain a single-cell suspension, the previous procedure for preparing brain sections was followed, replacing the isolation NMDG buffer with an isolation solution containing (in mM): 136.0 NaCl, 5.4 KCl, 10 4-(2-hydroxyethyl)-1-piperazineethanesulfonic acid (HEPES), 5.5 glucose, with pH of 7.4 and osmolality of 290 ± 3 mOsmol/kg. The tissue was cut into 500-700 µm coronal sections. Then, the cortex of the hemisphere with the ischemic injury (and the corresponding hemisphere in sham-operated animals) was excised and cut into smaller pieces with a razor blade. Subsequently, the cortical tissue was incubated with constant shaking at 37°C for 40 minutes in 1 ml of papain (20 U/ml) dissolved in the isolating solution supplemented with 60 µl of DNase (both from Worthington, Lakewood, NJ, USA). During the enzymatic dissociation, the tissue was gently triturated twice with a cut-open 1 ml pipette to break it up mechanically. After the dissociation procedure, the cell suspension was triturated with a 1-ml pipette and centrifuged at 300 × g for 3 min. The supernatant was then removed and the cells were resuspended in 0.9 ml of the isolation solution containing 100 µl of an ovomucoid inhibitor solution (both Worthington, Lakewood, NJ, USA) and 50 µl of DNase. Centrifugation at 300 × g for 3 min followed, the supernatant was removed, and the cell pellet was resuspended in Dulbecco’s phosphate-buffered saline (DPBS; Sigma-Aldrich, St. Louis, MO, USA) and filtered through a 70-µm pre-separation filter (Miltenyi Biotec, Bergisch Gladbach, Germany). The single-cell suspension was processed with a debris removal solution (Miltenyi Biotec) according to the manufacturer’s protocol. The resulting cell suspension was resuspended in Neurobasal-A medium (Life Technologies, Waltham, MA, USA) and stored at 4 °C for further processing.

### Fluorescence-activated cell sorting (FACS) and reverse-transcription quantitative polymerase chain reaction (RT-qPCR)

The single-cell suspension was stained at 4°C for 15 minutes with antibodies recognizing lymphocytes and endothelial cells (anti-CD45/CD31; BioLegend Way, San Diego, CA, USA) and labeling live cells (CellTrace calcein green; ThermoFisher Scientific, Waltham, MA, USA). All antibodies were diluted in Neurobasal-A containing 2% B27 (Life Technologies, Waltham, MA, USA). After staining, cells were washed in Neurobasal-A medium and stored at 4°C until FACS processing. Viable (Hoechst 33258-negative (Life Technologies, Carlsbad, CA, USA) and CellTracecalcein-green-positive) and CD45/CD31-negative cells were isolated by FACS (BD Influx, San Jose, CA, USA). The selected cells were either sorted into 1.5 ml Eppendorf tubes containing 200 µl of Neurobasal-A with 2% B27 and analyzed using a 10x Genomics single-cell RNA sequencing approach, or used for RNA isolation using the RNeasy Micro Kit (Qiagen, Valencia, CA, USA) according to the manufacturer’s protocol. The cDNA synthesis was performed with random hexamers. For reverse transcription of RNA, MAXIMA reverse transcriptase (ThermoFisher Scientific) was used. Quantitative RT-PCR was performed in technical triplicates using SYBR Green I Master Mix in a LightCycler 480 instrument (both from Roche Diagnostics, Indianapolis, IN, USA). Primers for RT-qPCR are listed in Supplementary Table S1.

### Single-cell RNA sequencing (scRNA-seq)

The Chromium Controller (10x Genomics, Pleasanton, CA, USA) was used to generate barcoded single-cell cDNA libraries using the Chromium Next Gem Single Cell 3’ kit, v3.1 following the manufacturer’s protocol. The barcoded cDNA was then pooled and sequenced using the NextSeq 500 sequencer (Illumina, San Diego, CA, USA) yielding over 100,000 reads per cell. The acquired sequencing data were then analyzed in R Studio (R-tools Technology Inc., Richmond Hills, Ontario, Canada) using Seurat (Stuart et al. 2019). All datasets were merged into a combined dataset of 12,850 cells and cell clusters were identified using Louvain approach based on principal component analysis (PCA). Nonlinear dimensionality reduction by Uniform Manifold Approximation and Projection (UMAP) (Becht et al. 2018) was applied to visualize low-dimensional embedding of the data confirming cluster assignment of cells. The obtained cell clusters were further analyzed based on specifically expressed genes previously reported in a molecular atlas of adult brain cells (Mizrak et al. 2019). Data compatible with the minimum information for a microarray experiment (MIAME) were deposited in the ArrayExpress repository within the EMBL-EBI BioStudies database under accession number E-MTAB-11967.

### Patch-clamp recording

Membrane properties of NG2 glia were recorded 3 days after MCAO; tdTomato-positive oligodendrocytes and perivascular cells were determined by their electrophysiological properties and were not analyzed. The patch-clamp technique in the whole-cell configuration, described in detail in our previous study (Kriska et al. 2021), was used to assess the membrane properties of NG2 glia and their progeny. Pre-incubated tissue sections were placed at room temperature in the recording chamber perfused with aCSF and gassed with 5% CO2. Recording pipettes with a tip resistance of 8-12 MΩ were made of borosilicate capillaries (Sutter Instruments, Novato, CA, USA) and filled with intracellular solution containing (in mM): 130 KCl, 0.5 CaCl_2_, 2 MgCl_2_, 5 EGTA, 10 HEPES (pH 7.2). Electrophysiological data were measured at a sampling frequency of 10 kHz with an EPC9 or an EPC10 amplifier, controlled by PatchMaster software (HEKA Elektronik, Lambrecht/Pfalz, Germany), and filtered with a Bessel filter. The resting membrane potential (V_M_) was measured by switching the amplifier to the current-clamp mode. FitMaster software (HEKA Elektronik, Lambrecht/Pfalz, Germany) was used to calculate the input resistance (IR) from the current value 40 ms after the onset of the depolarizing 10 mV pulse from the holding potential of −70 mV. Membrane capacitance (C_M_) was automatically determined by the software from the lock-in protocol. Current patterns were obtained by hyperpolarizing and depolarizing the cell membrane from the holding potential of −70 mV to values between −160 mV and 40 mV at 10 mV intervals. The pulse duration was 50 ms. To isolate the delayed outwardly rectified (K_DR_) and the inwardly rectified (K_IR_) potassium (K^+^) current components, a voltage step from −70 to −60 mV was used to subtract the time- and voltage-independent currents. To activate only the K_DR_ currents, cells were depolarized from the holding potential to −50 mV, and the amplitude of the K_DR_ currents was measured at 40 mV, 40 ms after the onset of the pulse. K_IR_ currents were determined analogously at −140 mV, also 40 ms after the onset of the pulse, while cells were held at - 70 mV. The fast-activating, fast-inactivating, outwardly rectified currents of the A-type K^+^ channel (K_A_) were measured at 40 mV and determined by subtracting the current traces with a hyperpolarizing prepulse of −110 mV from those with a depolarizing prepulse of −50 mV, and their amplitude was measured at the peak value. Current densities were calculated by dividing the maximum current amplitudes by the corresponding C_M_ values for each cell.

### Immunohistochemistry and confocal microscopy

Immunofluorescence staining was performed according to a previously described protocol (Kriska et al. 2021). The following primary antibodies were used: anti-APC mouse monoclonal antibody (1:200; Merck Millipore, Burlington, MA, USA), anti-GFAP-488 mouse monoclonal antibody (1:300; Thermo Fisher Scientific), anti-Mcm2 rabbit monoclonal antibody (1:800, Cell Signaling Technology, Danvers, MA, USA), anti-PCNA mouse monoclonal antibody (1:800; Abcam, Cambridge, UK), anti-Pcp4 rabbit polyclonal antibody (1:1.000; Thermo Fisher Scientific), anti-Mag mouse monoclonal antibody (1:800; Merck), anti-NeuN mouse monoclonal antibody (1:200; Merck), anti-Pdgfrα rabbit polyclonal antibody (1:200; Santa Cruz Biotechnology, Dallas, TX, USA), anti-vimentin rabbit monoclonal antibody (1:200, Cell Signaling Technology, Danvers, MA, USA). Secondary antibodies were donkey/goat polyclonal anti-rabbit/mouse IgG conjugated to Alexa-488 (1:200; Thermo Fisher Scientific). Counterstaining of nuclei was performed with 300 nM 40,6-diamidino-2-phenylindole (DAPI; Thermo Fisher Scientific).

Localization of cell types observed throughout the cortical area around the ischemic injury was assessed using a Dragonfly confocal fluorescence microscope with a 530 Andor rotating disk (Oxford Instruments, Oxford, UK) equipped with a Zyla 4.2 PLUS sCMOS camera and fusion acquisition system. Tile images were obtained by superimposing 50-70 single confocal images acquired with a 20× objective. The image series were then digitally fused using the fusion stitching tool. The acquired images were processed using Imaris visualization software (Oxford Instruments). Colocalization of the antibody immunofluorescence signal and endogenous fluorescence of NG2 glia was examined using the Leica Stellaris confocal platform (Leica Microsystems, Wetzlar, Germany). The immunopositive areas of the cortex surrounding the ischemic injury were further quantified using Fiji ImageJ software (NIH, Bethesda, MD, USA) (Schneider, Rasband, and Eliceiri 2012). Areas corresponding to cells with endogenous NG2-tdTomato fluorescence were calculated using the IJM script language written for the Fiji software package (fiji.sc) and normalized to the DAPI-positive area. First, the confocal data were split into individual channels. Then, the DAPI-stained nuclei were segmented using the threshold dialog, and the contiguous segmented nuclei were subdivided using the watershed method. The number of nuclei was then counted using the Particle Analyzer Tool in Fiji. A similar thresholding approach for segmentation was also applied to the antibody signal. Finally, we identified and counted individual NG2-tdTomato^+^ cells that had positive antibody signal using the Image Calculator in Fiji, which can generate an image with the intersections of two images, and the Particle Analyzer, which can count the number of overlaps in the image. For each quantification, a total of six cortical sections from at least two biological replicates were evaluated.

### Data analysis and statistics

Cells measured by the patch-clamp technique were divided into four groups based on their electrophysiological properties. The electrophysiological properties were then compared with the Kruskal-Wallis post-test using the one-way ANOVA method. Data are presented as means ± standard error of the mean (S.E.M.) or means ± standard deviation (S.D.) for a given number (n) of samples/cells. ANOVA was performed to detect significant differences among experimental groups in immunofluorescence quantification and RT-qPCR. Values of p < 0.05 were considered significant.

## RESULTS

### Ischemic brain injury results in increased numbers of NG2-expressing cells and their accumulation at the margin of the lesion

To investigate the nature of NG2-expressing glial cells in the healthy and ischemic cortex of adult rodents, we used transgenic mice that allow visualization of NG2 glia by tamoxifen-mediated Cre recombination. We used mice that had the gene encoding the tandem dimer (td) of the red fluorescent protein variant ‘tomato’ integrated downstream of the Rosa26 locus (Madisen et al. 2010). These “reporter mice” were crossed with mice possessing a Cre-ERT2 cassette encoding the tamoxifen-inducible Cre recombinase fused to estrogen receptor T2 under the control of the regulatory region of the Cspg4/NG2 gene (Zhu et al. 2011). In the resulting mouse strain, R26-tdTomato/Cspg4-CreERT2, tamoxifen-inducible excision of a transcriptional blocker located upstream of the tdTomato cDNA results in red fluorescence in Cspg4/NG2-expressing cells and their progeny (Fig. 1A).

**Figure 1.**
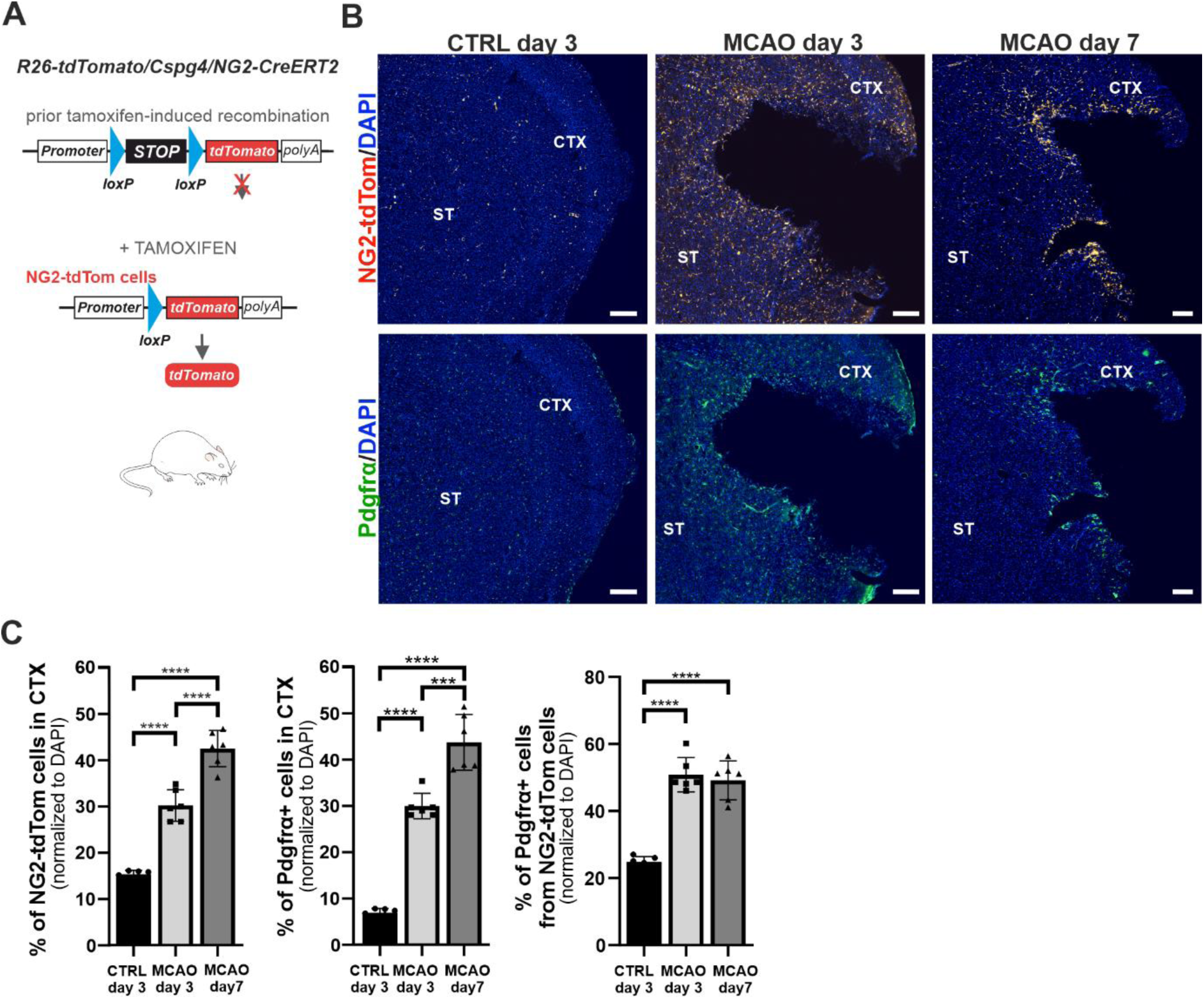
Neural-glial antigen 2 (NG2)-positive cells (or their progeny) surround the wound in the adult cortex 7 days after ischemic injury. (A) Schematic representation of the mouse model for tamoxifen-induced production of red fluorescent protein tdTomato in NG2-expressing cells. (B) Fluorescence micrographs of the coronal section of the left hemisphere of sham-operated (CTRL) mice and mice that underwent middle cerebral artery occlusion (MCAO) 3 or 7 days after surgery. The upper images show cells labelled with the red fluorescent protein tdTomato (NG2-tdTom; red fluorescent signal), and the lower images show cells stained with the antibody against platelet-derived growth factor receptor alpha (Pdgfrα) (green fluorescent signal). Samples were counterstained with 4’,6-diamidino-2-phenylindole (DAPI) nuclear stain; scale corresponds to 0.15 mm. (C) Quantification of cells with red (left graph) or green (middle) fluorescence in the cortex 3 and 7 days after MCAO compared with CTRL mice. The percentage of cells was determined in relation to DAPI-positive cells in the brain area examined. The percentage of Pdgfrα-positive cells to tdTomato-positive cells is shown in the right graph. At least six samples from two littermates (n ≥ 6) in each group were analyzed. Error bars represent SD. Statistical significance was determined using a one-way ANOVA test; *** p < 0.001; **** p < 0.0001. CTX, cortex; ST, striatum.

The Cre-loxP system was activated two weeks before MCAO with tamoxifen, and sham-operated mice (CTRL) were used as controls. Native red fluorescence of tdTomato protein activated after tamoxifen treatment in cells expressing the CSPG4-CreER2 transgene was used as a surrogate marker for NG2 glia and their progeny. These “red” cells, termed NG2-tdTom, were significantly more numerous in the cerebral cortex after ischemia compared with control tissue. Moreover, NG2-tdTom cells formed a rim around the injury site 7 days after MCAO, either because of their migration to the injury site or because of a strong proliferative response of cells in the injured area. The transmembrane proteoglycan NG2 regulates progenitor cell migration in response to injury and its polarization toward the acute lesion (reviewed in (Biname 2014)), but NG2 expression in the CNS is not restricted to glial cells. Recent studies have uncovered a subtype of perivascular cells whose markers include platelet-derived growth factor receptor beta (Pdgfrβ) and NG2. These perivascular cells exhibit mesenchymal stem-cell properties and are involved in spinal cord injury repair (Zhu et al. 2022). Nevertheless, NG2 glia are also characterized by Pdgfrα expression (reviewed in (Nishiyama et al. 2016)), and therefore we used immunodetection of Pdgfrα to distinguish these cells from NG2^+^ perivascular cells. Pdgfrα^+^ cells followed the increasing amount of NG2-tdTom cells lining the ischemic lesion after MCAO, but the proportion of detected Pdgfrα^+^ cells remained approximately the same at 3 and 7 days after injury (Fig. 1B,C). To gain deeper insight into the cellular changes caused by the brain injury, we performed gene expression profiling of NG2-tdTom cells from the ischemic cortex and compared them with cells from the cortex of sham-operated mice. Live cells were isolated by FACS based on endogenous tdTomato fluorescence and cell viability marker CellTrace calcein green-488; vascular endothelial cells and leukocytes, i.e., CD31/CD45-positive cells, were excluded from sorting. Remarkably, we observed distinct populations of NG2-tdTom cells based on fluorescence intensity (Suppl. Fig. S1A). We hypothesize that the less intense red fluorescence is related to active proliferation of NG2 glia (Psachoulia et al. 2009) and that the cells with more intense fluorescence include their derivatives and NG2^+^ perivascular cells. Therefore, we subsequently examined the expression of tdTomato and markers of the major cell types in the cortex (Zeisel et al. 2015), which confirmed the predominant presence of NG2 glia, oligodendrocytes, and perivascular cells within the sorted NG2-tdTom cells (Suppl. Fig. S1B).

Single-cell RNA-seq analysis was performed on isolated NG2-tdTom cells 3 days after MCAO. To cover a larger time span and obtain derivatives of NG2^+^ cells, NG2-tdTom cells were also analyzed 12 days after MCAO. Corresponding control cells from sham-operated animals were obtained at both time points. The sequences obtained were analyzed using the Seurat package (Stuart et al. 2019) for comprehensive integration of scRNA-seq data. A merged dataset containing all MCAO and CTRL samples was used to determine cell clusters that were assigned to three expected cell types based on specifically expressed genes (Mizrak et al. 2019). We identified NG2 glia expressing the *Pdgfrα* and tenascin R (*Tnr*) genes and three clusters of oligodendrocytes with myelin-associated glycoprotein (*Mag*) and myelin-oligodendrocyte glycoprotein (*Mog*) mRNA. The remaining 10 clusters were assigned to perivascular cells and their derivatives, based on the expression of markers *Pdgfrβ* and regulator of G protein signaling 5 (*Rgs5*) (Fig. 2A). Genes with significantly increased expression in the corresponding cluster compared to all other cells in the sample can be found in Suppl. Table S2. Among the isolated NG2-tdTom cells, pericyte-related cells were the most abundant. The total number of these cells varied slightly depending on whether CTRL or post-ischemic brain tissue was analyzed. However, because the focus of our study was on NG2 glia and their derivatives, we excluded all pericyte clusters from further analyses. We detected a higher proportion of NG2 glia in the ischemic cortex; moreover, their number decreased less rapidly on day 12 in the ischemic sample than in the healthy brain. In contrast, the number of mature oligodendrocytes decreased more rapidly after MCAO (Fig. 2B). Oligodendrocyte transcription factor 2 (*Olig2*) expression was present throughout the oligodendroglial lineage, with similar levels in ischemic and CTRL tissues. *Olig2* expression was highest in NG2 glia (cluster 0) and gradually decreased in the three (1-3) oligodendrocyte (OL) clusters (Fig. 2C). In addition, the differential degree of OL maturation was illustrated by the decreasing expression of the glutamate ionotropic receptor AMPA-type subunit 2 (*Gria2*) and the increased expression of the K^+^ and Na^+^ conducting hyperpolarization-activated cyclic nucleotide-gated ion channel 2 (*Hcn2*) genes (Suppl. Fig. S2) in clusters 1 to 3, as reported by Kirdajova and colleagues (Kirdajova et al. 2021). Interestingly, the OL maturation process was not altered in the ischemic tissue, so the lower number of OL after MCAO can be explained by their death.

**Figure 2.**
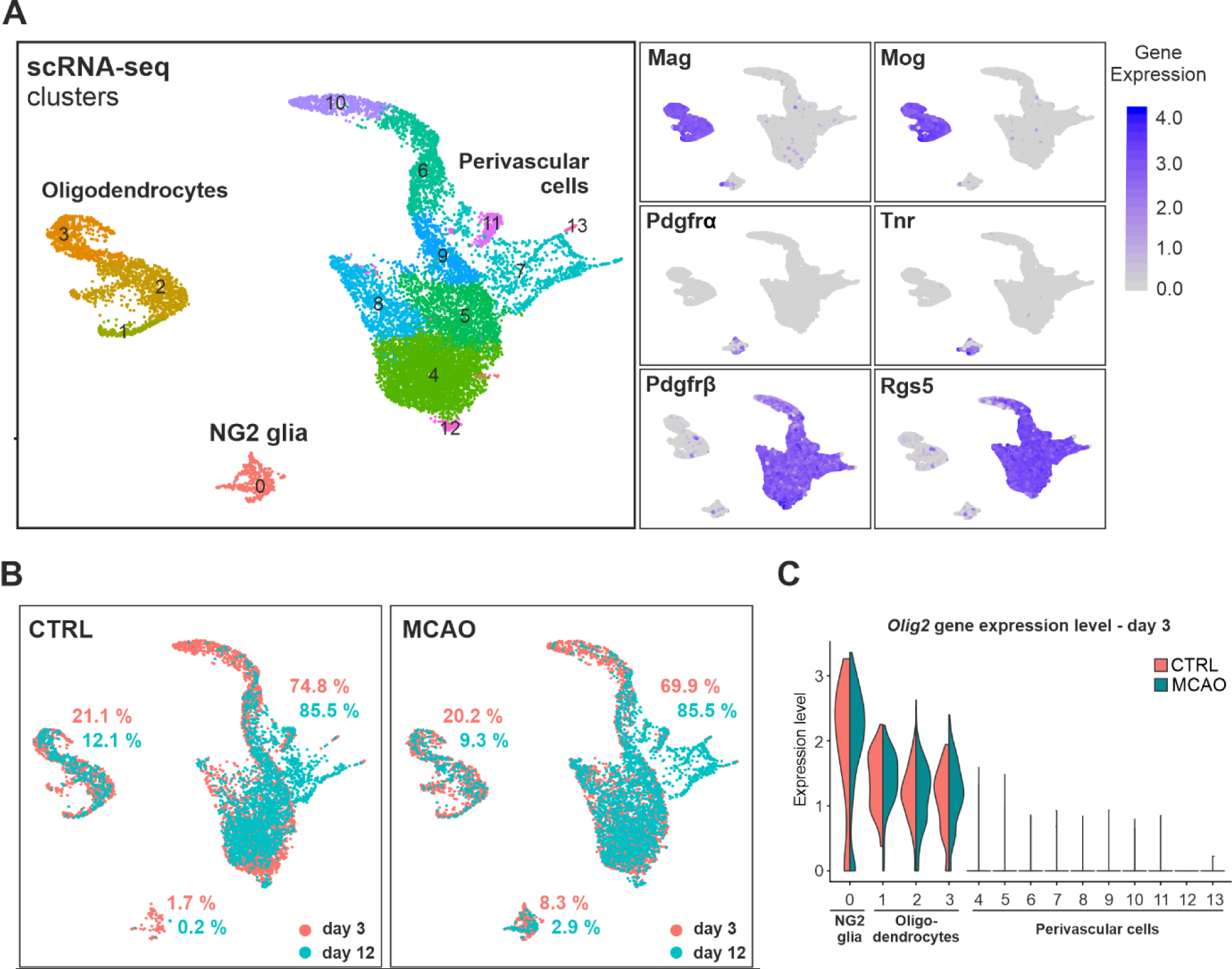
Single-cell RNA sequencing (scRNA-seq) of cells from the ischemic cortex and healthy tissue of adult mice. After the removal of apoptotic cells, we analyzed a pooled dataset consisting of 3,444 (day 3) and 2,919 (day 12) cells in the ischemic (MCAO) samples and 3,829 (day 3) and 2,658 (day 12) cells in the control tissue (CTRL) with an average transcript number of 2,743 per cell. (A) Left, three main cell types were classified into 14 different clusters based on principal component analysis (PCA) of the merged dataset and visualized by Uniform Manifold Approximation and Projection (UMAP). Right, feature plots show the expression level (indicated by the color scale) of genes specific to the corresponding cell type. (B) UMAP plots show the density of cells in each cluster of CTRL and MCAO samples in the two time points. The percentage of cells in the different cell populations is indicated. (C) Violin plot showing oligodendrocyte transcription factor 2 (*Olig2*) expression in the indicated cell clusters at day 3. Mag, myelin-associated glycoprotein; Mog, myelin-oligodendrocyte glycoprotein; Pdgfrα/β, platelet-derived growth factor receptor alpha/beta; Rgs5, regulator of G protein signaling 5; Tnr, tenascin R.

Despite the marked decrease in the number of mature OLs in the MCAO samples, we detected an increased number of newly developing OLs near the ischemic injury that were positive for adenomatous polyposis coli (APC) protein (Fig. 3A). APC is produced during OL maturation (Lang et al. 2013) and reflects remyelination of the ischemic cortex. As expected, the number of newly formed OLs increased rapidly after induction of FCI (Fig. 3B) and formed a lining along the ischemic scar at day 7. In addition, colocalization of red fluorescence with MAG protein indicated newly formed OLs derived from NG2-tdTom OPCs (Fig. 3A). The newly maturing OLs likely complement the number of mature OLs that die in the vicinity of the lesion after ischemic injury. This is also consistent with the fact that we observed only a small decrease in the number of mature OLs 3 days after MCAO.

**Figure 3.**
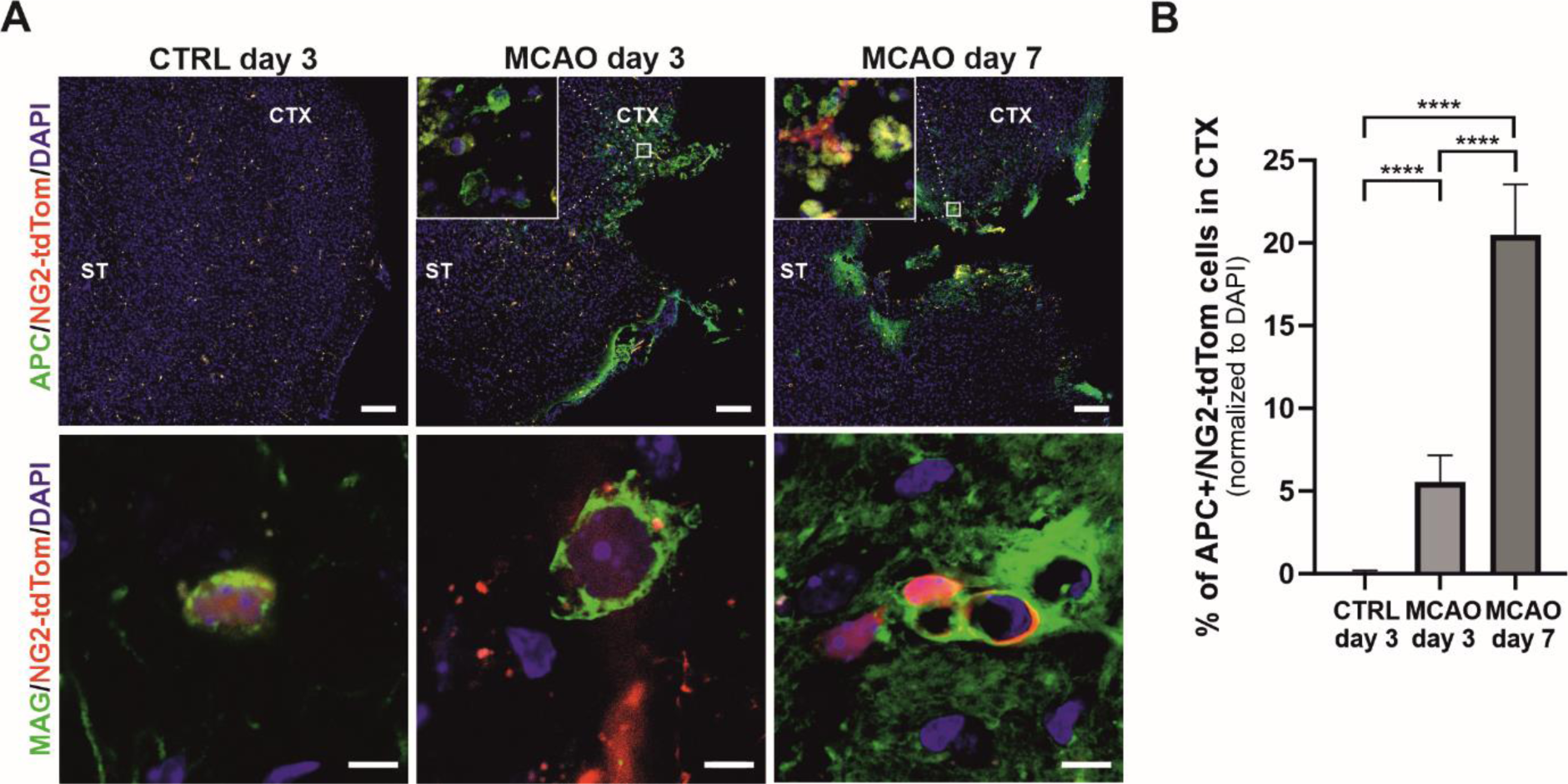
Differentiation of oligodendrocytes (OLs) from NG2 glial cells. (A) Representative slides show staining of NG2-tdTom cells and newly mature OLs (labeled with adenomatous polyposis coli (APC)) around the wound (upper panel; scale bar corresponds to 150 μm) and colocalization of the OL marker MAG with NG2-tdTom cells in the cortex (lower panel; scale bar corresponds to 5 μm) of the healthy and ischemic brains. Samples were counterstained with DAPI nuclear stain. Enlarged images are shown in the insets. (B) Quantification of APC/NG2-tdTom cells in the cortex surrounding the wound. Statistical significance was determined by one-way ANOVA test; **** p < 0.0001. CTX, cortex; ST, striatum.

Overall, after activation of the tdTomato reporter allele in NG2^+^ cells 2 weeks before FCI, we identified three major cell types in both ischemic and healthy tissue, namely NG2 glia, OLs, and perivascular cells. NG2 glia were significantly enriched 3 and 7 days after ischemic injury, whereas 12 days later, progressive loss of NG2 glia was already observed in the cortex. The loss of mature OLs 3 and 12 days after surgery was moderate after ischemia, and conversely, new OLs were formed with increasing frequency 3 and 7 days after injury.

### Five distinct subpopulations of NG2-tdTom cells expressing Pdgfrα were distinguished in the adult cortex

In recent publications, we have demonstrated the existence of different types of NG2 glia, including a newly defined population of astrocyte-like NG2 glia, in the CNS after different types of injury (Kirdajova et al. 2021, Valny et al. 2018). To further characterize NG2 glia and their gene expression in the adult brain regenerating after FCI, we used the scRNA-seq data and immunohistochemistry to analyze a subset of NG2-tdTom/Pdgfrα^+^ cells. As expected, the amount of NG2-tdTom/Pdgfrα^+^ cells in the sham-operated mice was relatively low 3 days after surgery, whereas we could not detect these cells at all in the tissue sample after 12 days. However, histological examination of brain tissue 7 days after MCAO showed that the number of NG2-tdTom/Pdgfrα^+^ cells increased, but by day 12, their number was already significantly lower (Fig. 4A). Therefore, we used only the samples isolated at day 3 for the analysis of the NG2-tdTom/Pdgfrα^+^ cell population. Analysis with the R package Seurat revealed five different clusters of NG2-tdTom/Pdgfrα^+^ cells. Interestingly, their frequency differed significantly in the sham-operated and ischemic tissues (Fig. 4B). To assign the obtained cell clusters to the already known subpopulations of NG2 glia, we determined the expression profile of each cluster in more detail (Suppl. Table S3).

**Figure 4.**
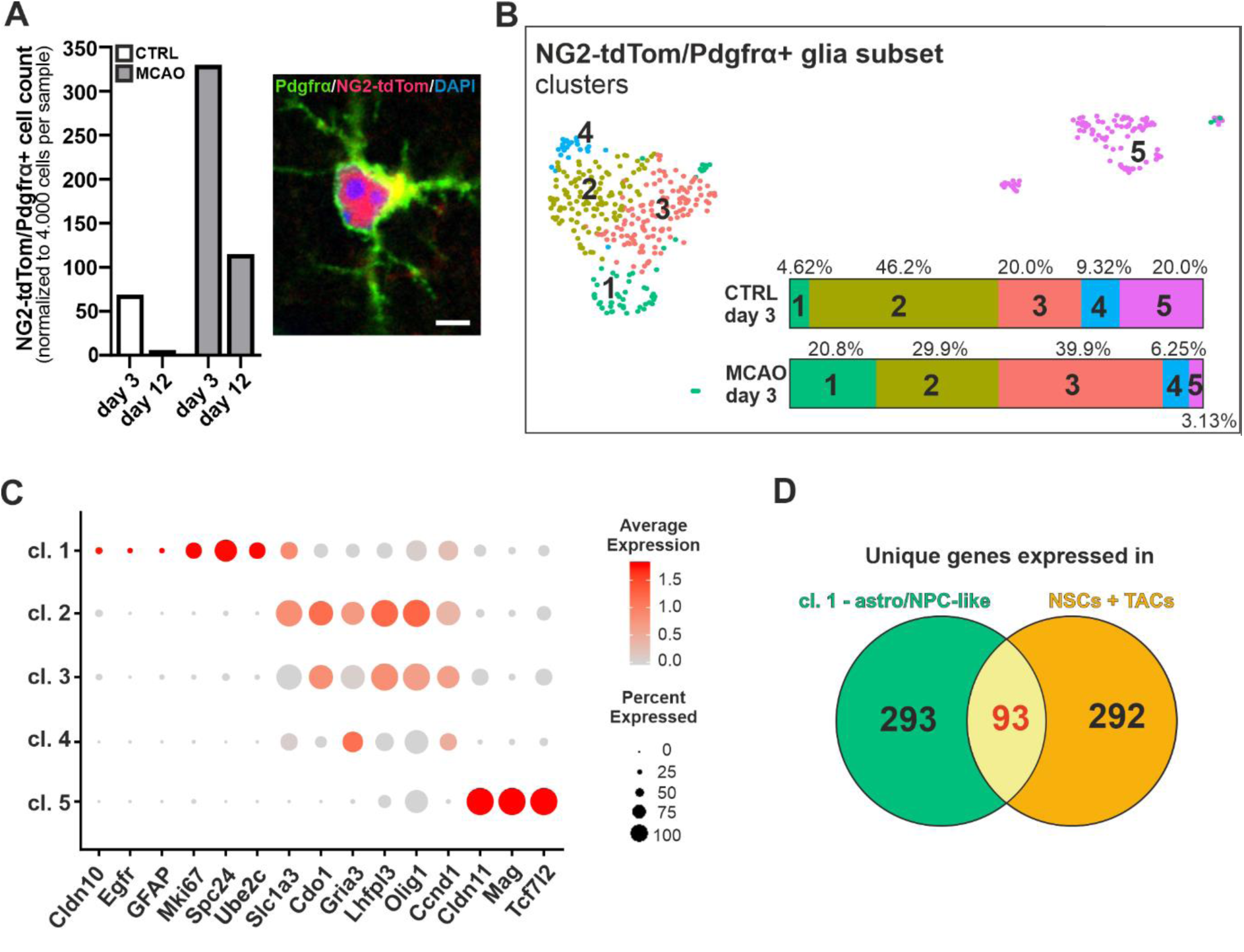
NG2-tdTom/Pdgfrα^+^ cell subpopulations and their gene expression. (A) Left, the number of NG2 glia in the ischemic and sham-operated cortex. The number of NG2-tdTom/Pdgfrα^+^ cells was normalized to 4000 cells per sample analyzed by scRNA-seq. Right, a representative image showing colocalization of anti-Pdgfrα staining and endogenous tdTomato fluorescence. The specimen was obtained from the cortex 3 days after the sham operation. The scale is 5 µm. (B) UMAP plot showing the distribution of NG2-tdTom/Pdgfrα^+^ cells in five identified clusters. On the bottom right the percentage of NG2 glia in each cluster in the indicated sample is shown; color coding corresponds to the UMAP plot. (C) Dot plot showing expression of selected genes in NG2-tdTom/Pdgfrα^+^ cells clusters identified by scRNA-seq. Color coding shows the change in gene expression compared with the average gene expression in all clusters; dot size corresponds to the percentage of cells expressing the selected gene within the cluster. (D) Venn diagram showing the number of genes specifically expressed in cluster 1, identified as the astro/NPC-like NG2 glia, and in the neural stem cells (NSCs) and transit amplifying cells (TACs) from the subventricular zone (SVZ) and striatum of the healthy brain (Kriska et al. 2021). The overlap between the two groups of genes is highlighted in red and the genes are listed in Suppl. Table S4. Ccnd1, cyclin D1; Cdo1, cysteine dioxygenase type 1; Cldn10/11, claudin 10/11; Egfr, epidermal growth factor receptor; GFAP, glial fibrillary acidic protein; Gria3, glutamate ionotropic receptor AMPA-type subunit 3; Lhfpl3, lipoma HMGIC fusion partner-like 3; Mki67, proliferation marker Ki-67; Slc1a3, solute carrier family 1 member 3; Spc24, SPC24 component of NDC80 kinetochore complex; Tcf7l2, transcription factor 7 like 2; Ube2c, ubiquitin conjugating enzyme E2 C.

Groups of genes characteristic of each cluster were analyzed by Gene Set Enrichment Analysis (GSEA) using the online tool Enrichr (Chen et al. 2013, Kuleshov et al. 2016, Xie et al. 2021). For each cluster, we identified the most important term related to the cell type (Cell marker augmented 2021 (Zhang et al. 2019)), cellular compartment (Jensen compartments (Binder et al. 2014)), biological process (gene ontology (GO) biological processes 2021 (Ashburner et al. 2000, Gene Ontology 2021)), and signaling pathway (WikiPathway 2021 human (Martens et al. 2021)), in which the acquired genes were mostly enriched (Table 1).

**Table 1.** Gene set enrichment analysis of NG2-tdTom/Pdgfrα^+^ cell clusters. Genes specifically expressed in each cluster (significance criterion: average expression |log2 FC | > 0.25 and p-value < 0.05) were analyzed using the indicated gene databases. The category with the most significant p-value (calculated from Fisher’s exact test; p-value < 0.0001) in each database is shown. Cluster coloring corresponds to the UMAP plot in Fig. 4B. Genes are listed in Suppl. Table S3.

| Cluster | Cell marker augmented 2021 | Jensen compartments | GO biological processes 2021 | WikiPathway 2021 human |
| --- | --- | --- | --- | --- |
| 1 | Neural progenitor cell: embryonic prefrontal cortex | Cytosolic ribosome | SRP-dependent cotranslational protein targeting to membrane (GO:0006614) | Cytoplasmic ribosomal proteins WP477 |
| 2 | Oligodendrocyte progenitor cell: embryonic prefrontal cortex | Synapse associated extracellular matrix | Nervous system development (GO:0007399) | Splicing factor NOVA regulated synaptic proteins WP4148 |
| 3 | Mitotic fetal germ cell: fetal gonad | Cytosolic ribosome | Cytoplasmic translation (GO:0002181) | Cytoplasmic ribosomal proteins WP477 |
| 4 | Cardiomyocyte: undefined | Axon initial segment | mRNA splicing, via spliceosome (GO:0000398) | mRNA processing WP411 |
| 5 | Oligodendrocyte: brain | Myelin sheath adaxonal region | Axon development (GO:0061564) | Glial cell different. WP2276 |

This correlated well with the expression of selected markers of OPCs sampled from the molecular atlas of adult brain cells reported previously (Mizrak et al. 2019). The expression of these genes, such as *Olig1*, cysteine dioxygenase type 1 (*Cdo1*), lipoma HMGIC Fusion Partner-Like 3 (*Lhfpl3*), and solute carrier family 1 member 3 (*Slc1a3*), was highest in cluster 2 of NG2 glia, which was most consistent with OPCs according to the GSEA, and gradually decreased in clusters 3 and 4 (Fig. 4C). To compare these clusters with the previously defined subpopulations of NG2 glia, we used the same set of cluster-specific genes for expression analysis as in the reference (Kirdajova et al. 2021) and found an overlap between our clusters 2, 3, and 4 and the two subpopulations of NG2 glia previously identified in the cerebral cortex after different types of injury (Suppl. Fig. S3A). These cell clusters are likely OPCs, and their differentiation into subpopulations is probably related to different stages of their life cycle. While cluster 2 most closely corresponded to basal OPCs, cluster 3 represented mitotic cells according to GSEA and additionally showed increased expression of *Dcx* and nestin. While the former gene is associated with cell migration in this context and is expressed in most OPCs (Boulanger and Messier 2017a), nestin expression was observed in dividing NG2 glia migrating to the edge of the lesion (Robins, Villemain, et al. 2013).

The ratio of these two groups changed in response to ischemic injury, with a decrease in basal OPCs and an increase in mitotic/migratory OPCs after MCAO. Cluster 4 contained the lowest number of OPCs and remained a mystery to us. Given the overall low gene expression, we speculate that these may be quiescent cells that form a pool of reserve OPCs that are activated after MCAO and therefore, they decrease in number in the ischemic cortex (Fig. 4B). Cluster 5 showed strong expression of genes associated with OL differentiation, such as claudin 11 (*Cldn11*), Mag, myelin basic protein (*Mbp*), or transcription factor 7 like 2 (*Tcf7l2*) (Fig. 4C and Suppl. Fig. S3A). *Cldn11* reflects the onset of OL differentiation and cell cycle arrest, whereas the *Mag* and *Mbp* genes represent markers of differentiation and their expression is further increased in myelinating OL (Yu et al. 2013). Interestingly, although *Tcf7l2* expression is responsive to Wnt signaling, the protein product of this gene, Tcf4, also functions as a transcriptional regulator of this signaling (Hrckulak et al. 2016) and its presence has been described in post-mitotic OPCs, where it coordinates OL maturation (Fu et al. 2009, Guo and Wang 2023, Zhao et al. 2016). Moreover, the frequency of the latter population of differentiating OLs decreased after MCAO (Fig. 4B), consistent with the previously demonstrated post-ischemic dynamics of NG2 glia subpopulations (Kirdajova et al. 2021).

In addition to the mitotic/migratory OPCs, cluster 1 was another population that was more numerous after MCAO than in the CTRL samples (Fig. 4B). In contrast to the gradual transition from OPCs to differentiated OLs observed in clusters 2-5, genes characteristic of OPCs or intermediate OPC-OLs were expressed minimally in cluster 1. Instead, we detected the expression of *Cldn10* and genes encoding the intermediate filaments GFAP, nestin, and vimentin. All of these genes are specific to reactive astrocytes or neural stem cells (NSCs) (de Pablo et al. 2013, Liu et al. 2014, Mignone et al. 2004, Mizrak et al. 2019, Wilhelmsson et al. 2019). We also observed epidermal growth factor receptor (*Egfr*) gene expression. The transmembrane protein encoded by this gene is critical for NSC activation (Cochard et al. 2021). Consistent with this, we also observed other genes characteristic of NSCs or transit amplifying cells (TACs), such as marker of proliferation Ki-67 (*Mki67*), NDC80 kinetochore complex component SPC24 (*Spc24*), and ubiquitin-conjugating enzyme E2 C (*Ube2c*) (Mizrak et al. 2019) (Fig. 4C and Suppl. Fig. S3B). Moreover, according to the most “enriched” term “Cell Marker Augmented 2021,” cluster 1-specific genes were most abundant in neural progenitor cells (NPCs; Table 1). Finally, we compared genes specific to this astro/NPC-like cluster (cluster 1) with genes whose expression was described by our group in NSCs and TACs isolated from the subventricular zone and striatum (SVZ+S) of adult mice (Kriska et al. 2021). These two sets of genes overlapped for about a quarter of the genes, i.e., 93 genes were found in both sets (Fig. 4D, genes are listed in Suppl. Table S4).

In summary, based on gene expression of characteristic markers, we characterized five NG2-tdTom/Pdgfrα^+^ cell subtypes as follows: (1) the oligodendroglial lineage, which includes basal OPCs, mitotic/migratory OPCs, putative quiescent OPCs, and transit OPCs that differentiate to OLs; (2) the previously identified astrocyte-like NG2 glial cells. In addition, we found that the astrocyte-like NG2 glia subpopulation shares a particular expression profile with NPCs isolated from the SVZ+S of the adult brain.

### Functional analysis of post-ischemic NG2-tdTom cells

As previously shown *in vitro*, cortical NG2 glia form several distinct subpopulations that differ in their electrophysiological properties and may represent cells with different cell fates (Honsa et al. 2016). Our goal was to functionally distinguish subtypes of NG2-tdTom cells using the patch-clamp technique in brain slices and assign them to the subpopulations identified by scRNA-seq. Electrophysiological recordings were performed on post-ischemic NG2-tdTom cells in cortical brain slices 3 days after MCAO; tdTomato-positive OLs and perivascular cells were identified by their morphology and electrophysiological properties and were not analyzed. Based on their electrophysiological properties, we determined four distinct groups of cells (Fig. 5). The most numerous group (n = 45) consisted of undifferentiated NG2 glia, previously described as “complex” cells (Kriska et al. 2021). These cells exhibited both K_IR_ currents and outwardly rectified voltage-dependent K^+^ currents represented by K_DR_ and K_A_ currents (Fig. 5A). The membrane properties (V_M_, IR, C_M_, and densities of K_IR_, K_DR_, and K_A_ currents) of the other three subpopulations were compared with this group of cells. Cells with a markedly depolarized V_M_, a very high IR, and a low density of K_IR_ currents were identified as migrating NG2 glia (Fig. 5B; n = 6), because cells with such electrophysiological properties had previously been identified as Dcx-positive migrating OPCs (Boulanger and Messier 2017a, Kriska et al. 2021).

**Figure 5.**
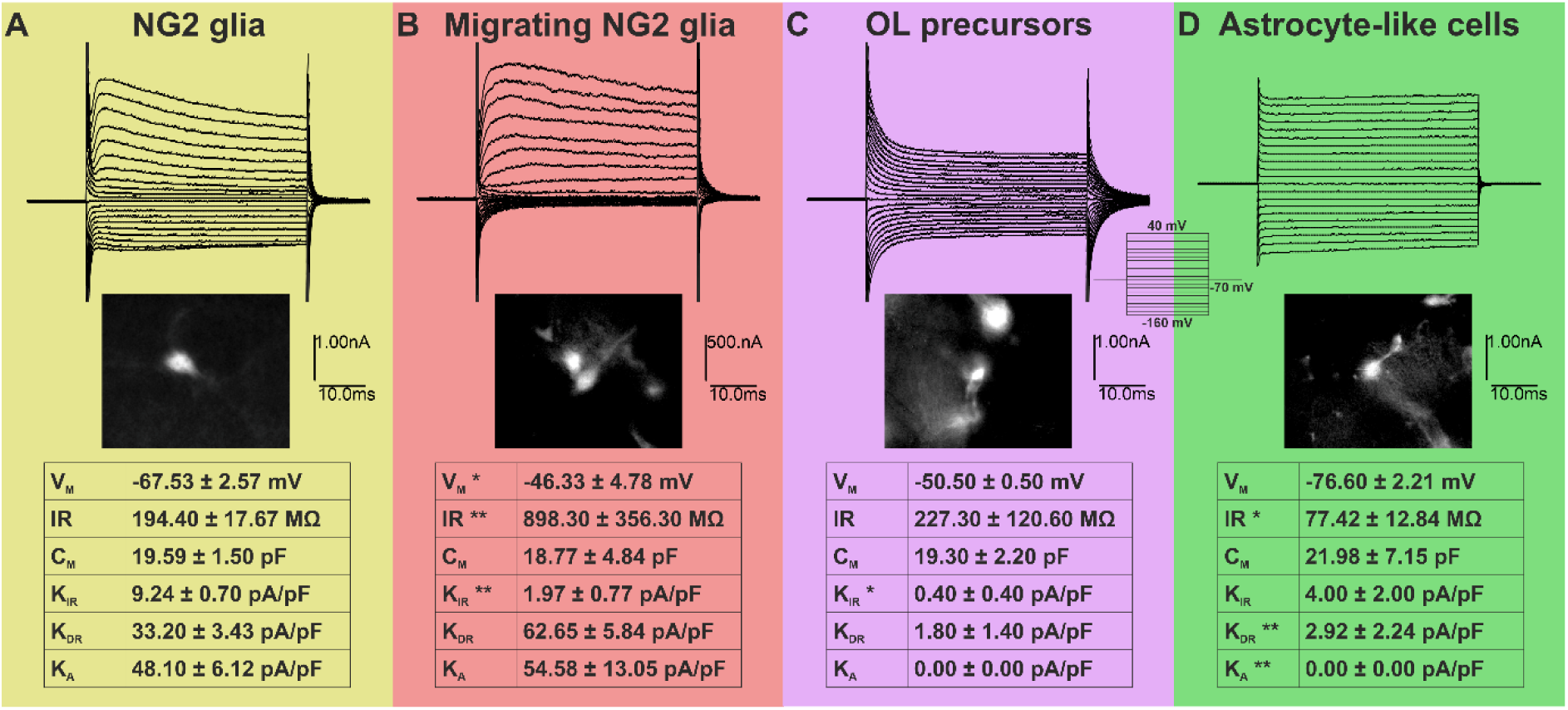
NG2-tdTom cells form electrophysiologically distinct subpopulations after ischemic injury. (A) NG2 glia in the progenitor state show both inwardly (K_IR_) and outwardly rectified K^+^ currents (delayed outwardly rectified K^+^ currents – K_DR_, A-type, transient K^+^ currents – K_A_) that are voltage-dependent; n = 45. (B) Migrating NG2 glia are characterized by reduced inward conductance, i.e., low density of K_IR_ currents. They also show a more depolarized resting membrane potential (V_M_) and higher input resistance (IR); n = 6. (C) NG2 glia differentiating into oligodendrocytes (OL) show voltage-independent decaying K^+^ currents, lower density of K_IR_ currents, and no K_A_ currents; n = 2. (D) NG2 glia with astrocyte-like, voltage-independent, nondecaying K^+^ currents show lower IR, lower current density of K_DR_, and no K_A_ currents; n = 5. Current responses/profiles were obtained by applying hyperpolarizing and depolarizing pulses ranging from −160 mV to 40 mV in 10 mV increments (see the inset). All current values in the table are stated as current densities (the current amplitude divided by the C_M_). Statistical significance was determined using the one-way ANOVA test and the Kruskal-Wallis post-test. Significant differences from values measured in NG2 glia (A) are indicated with asterisks; * p < 0.05; ** p < 0.01. C_M_, membrane capacitance; mV, millivolts; pA, picoampere; pF, picofarad.

In addition, white matter NG2 glia have been shown to lack K_IR_ currents (Chittajallu, Aguirre, and Gallo 2004, Diers-Fenger et al. 2001), leading us to hypothesize that these cells are white matter NG2 glia that have migrated into the injured ischemic cortex. Two cells with voltage-independent decaying currents were identified as NG2 glia that had already proceeded to OL differentiation (Fig. 5C). This subpopulation was characterized by a very low current density of K_IR_ compared to the “complex” cells. In addition, these cells lacked K_A_ currents and had a very low K_DR_ current density. These results were not statistically significant, but we believe that this is mainly due to the low abundance of this cell type in the sample. The last group was represented by cells with symmetric, voltage-independent, nondecaying currents (Fig. 5D). These cells (n = 5) were previously described as astrocytes (Honsa et al. 2016) and shared a common electrophysiological profile with astrocyte-like NG2 glia (Kirdajova et al. 2021). The large passive conductivity of these cells was represented by a low IR, the absence of K_A_ currents, and a low density of K_DR_ currents.

Overall, we identified four different types of cells based on their electrophysiological properties. We hypothesize that these cells correspond to the subpopulations identified by scRNA-seq. However, we were unable to identify a subpopulation of NG2 glia that is sparsely represented in the ischemic brain and that we designated as quiescent. This subpopulation was distinguished only by differential gene expression, so its electrophysiological characterization and function in the adult cortex is still unclear.

### Astro/NPC-like NG2 glia characterization

GFAP-positive astrocytes are the major type of glia that maintain homeostasis in the healthy brain. In brain injury, they are activated to limit CNS damage (reviewed in (Hol and Pekny 2015)). While GFAP^+^ cells are sporadic in the healthy adult cortex, significant proliferation and migration of these cells to the injury site occurs after FCI, peaking around day 7 after MCAO (Fig. 6A, B). Moreover, the shape and morphology of these cells are altered in reactive gliosis, as evidenced by an increase in the cell area stained with an anti-GFAP antibody (Fig. 6B). At the same time, NG2-tdTom/GFAP^+^ cells are detectable in the ischemic cortex as early as day 3 after MCAO (Fig. 6A). Although the highest increase in their number occurs at day 7 after the induction of FCI (Kirdajova et al. 2021), the expression of the gene encoding GFAP peaks as early as day 3 in NG2-tdTom cells and gradually decreases in the following days (Fig. 6C). Although we detected a small number of astro/NPC-like glia (cluster 1) in the CTRL tissue (Fig. 4B) based on their expression profile, GFAP expression was very low in these cells and increased markedly in the ischemic cortex (Fig. 6C and Suppl. Fig. S3B). In contrast, the expression of vimentin in NG2-tdTom/Pdgfrα^+^ cells was observed in the CTRL cortex (Suppl. Fig.S3B), and a portion of NG2-tdTom cells in the tissue showed concomitant staining for vimentin in the cortex of sham-operated mice, whereas these cells were more numerous in the ischemic cortex (Fig. 6A). These findings support our hypothesis that astro/NPC-like NG2 glia are present in small numbers in the healthy cortex but are not GFAP-positive; the appearance of GFAP expression in ischemic astro/NPC-like NG2 glia may thus reflect their activation in a manner similar to reactive astrogliosis.

**Figure 6.**
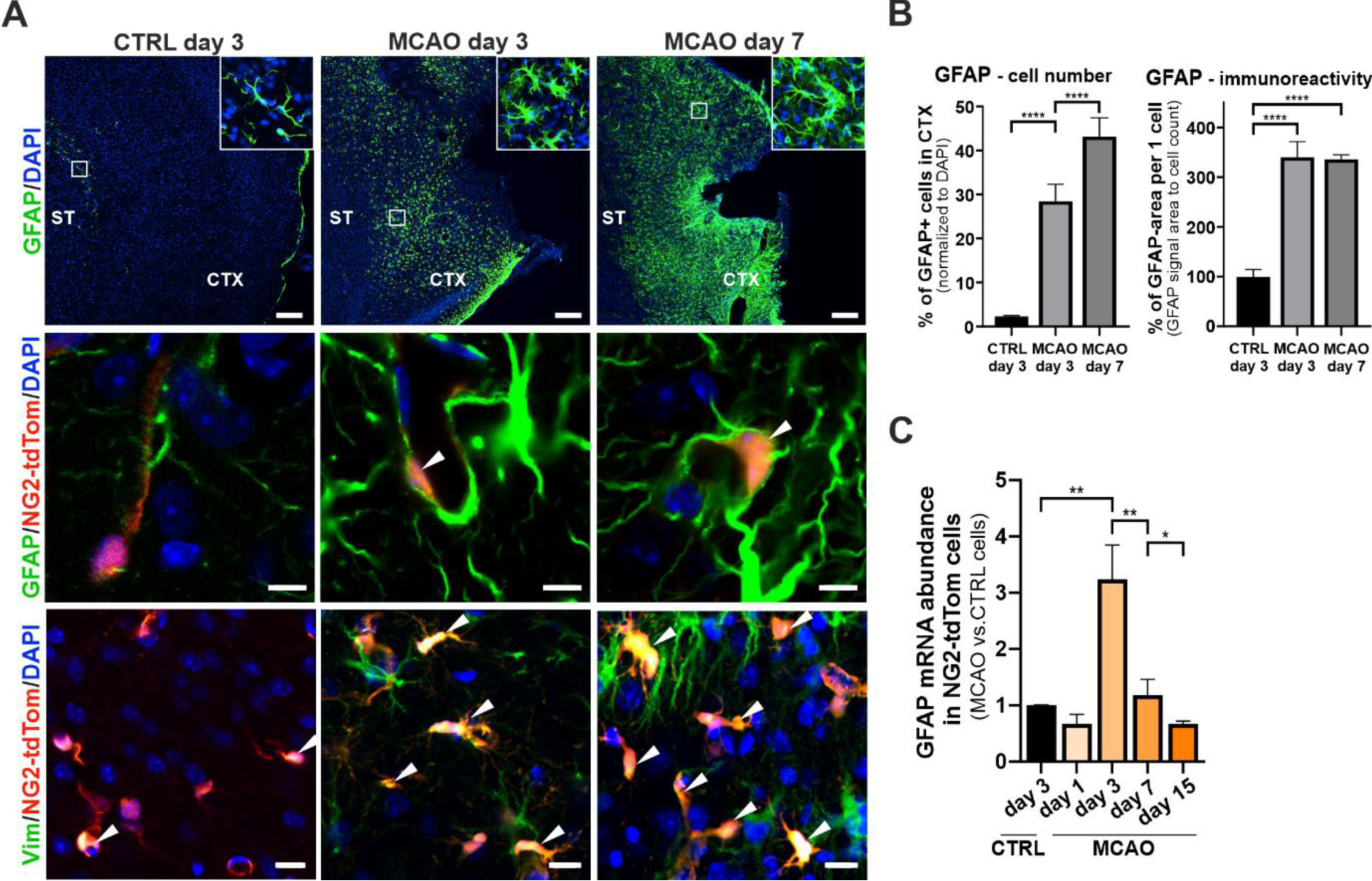
Detection of GFAP-positive cells in the ischemic brain. (A) Representative images showing GFAP^+^ astrocytes in the cortex surrounding the wound (upper panel; scale bar: 150 μm) and the presence of GFAP-positive NG2-tdTom cells (middle panel, white arrowheads; scale bar: 5 μm) and vimentin (Vim)-positive NG2-tdTom cells (lower panel, white arrowheads; scale bar: 10 μm) in the wound region. Enlarged images are shown in the insets. (B) Quantification of the number of GFAP^+^ cells (left) and the average area of the cell (right) in the ischemic and healthy cortex. (C) GFAP expression in the NG2-tdTom cells fluorescence-activated cell sorted at different days after MCAO was analyzed by reverse transcription quantitative polymerase chain reaction (RT-qPCR). Relative GFAP expression in the control sample (CTRL; analyzed on day 3 after sham surgery) was arbitrarily set to 1. Error bars represent SD of the technical triplicates. Statistical significance (panels B, C) was determined using a one-way ANOVA test; *, p < 0.05; **, p < 0.01; ****, p < 0.0001. CTX, cortex; ST, striatum.

In the healthy adult brain, GFAP and vimentin expression is restricted to astrocytes and the early stages of their transition to NSCs (Bernal and Peterson 2011, Mizrak et al. 2019). Moreover, we demonstrated in our recently published scRNA-seq analysis of cells from the SVZ+S that GFAP is a specific marker of NSC-transient astrocytes in the healthy brain tissue (Kriska et al. 2021). We used these scRNA-seq data to assign a number of additional markers to the neurogenic cell lineage and NG2 glia in healthy brain tissue (Suppl. Fig. S4A). Interestingly, GFAP expression was not detectable in NG2 glia, even after ischemia. This could be due to the low number of NPC-like cells in the NG2 glia population, which thus “escaped” detection when unsorted cells from SVZ+S tissue were used for scRNA-seq analysis. Another possibility is that these cells are mainly found in the cortex and not in the SVZ+S. In contrast, ischemic injury resulted in GFAP expression in all astrocyte populations, probably as a consequence of reactive gliosis. However, we found that several genes expressed in NPCs and mature neurons are also simultaneously present in NG2 glia and their expression appears to be upregulated after MCAO (Suppl. Fig. S4B). Therefore, we also analyzed the expression of these genes in the subpopulations of NG2-tdTom/Pdgfrα^+^ cells. Primarily, this was the achaete-scute family bHLH transcription factor 1 (*Ascl1*) gene, which encodes an activator of NSCs and induces formation of neurons and OLs (Matsubara, Matsuda, and Nakashima 2021); the expression of this factor was also observed in OPCs (Sueda and Kageyama 2021). After ischemic injury, the expression of *Ascl1* was decreased in the OPC subpopulation but increased in the astro/NPC-like subpopulation in the NG2-tdTom/Pdgfrα^+^ cells examined. In addition, we observed the expression of genes related to NSCs and TACs, such as minichromosome maintenance complex component 2 (*Mcm2*), *Mki67*, ribonucleotide reductase-regulating subunit M2 (*Rrm2*), and *Ube2c* (Suppl. Fig. S4B). These genes are functionally associated with cell division; in our scRNA-seq, their expression was restricted to the astro/NPC-like cluster (Fig. 7A). The Mcm2 protein is involved in the formation of the pre-replication complex that initiates genome replication. The assembly of this complex is essential during the G1 phase of the cell cycle to enable DNA replication in the subsequent S phase (Stoeber et al. 2001). The presence of Mcm2 in proliferating cells correlates well with Ki67 positivity, which has multiple functions during mitosis (Wharton et al. 2001). In the CTRL cortex, we detected a small amount (approximately 3%) of Mcm2-positive cells, and this amount was maintained at both time points examined. The number of Mcm2-positive cells increased significantly in the ischemic tissue 3 and 7 days after FCI induction (18.5% and 19.1%, respectively). However, we also observed a significant overlap of Mcm2 and tdTomato protein production in CTRL tissues. This overlap was similar at both time points (19.5% and 17.3%, respectively). After the ischemic injury, the overlap increased significantly to 48% and 38.9% at days 3 and 7 after MCAO, respectively. Of note, the number of Mcm2^+^ cells that were not positive for tdTomato increased at later time points after injury, likely reflecting proliferation of other cell types unrelated to NG2-tdTom cells (Fig. 7B,C). At the mRNA level, *Mcm2* gene expression in NG2-tdTom cells was highest on day 1 after FCI and gradually decreased over time. In contrast, the expression of the *Mki67* gene, which encodes a G2/mitotic marker (Uxa et al. 2021), was significantly increased only from day 3 after FCI and then decreased steeply (Fig. 7D).

**Figure 7.**
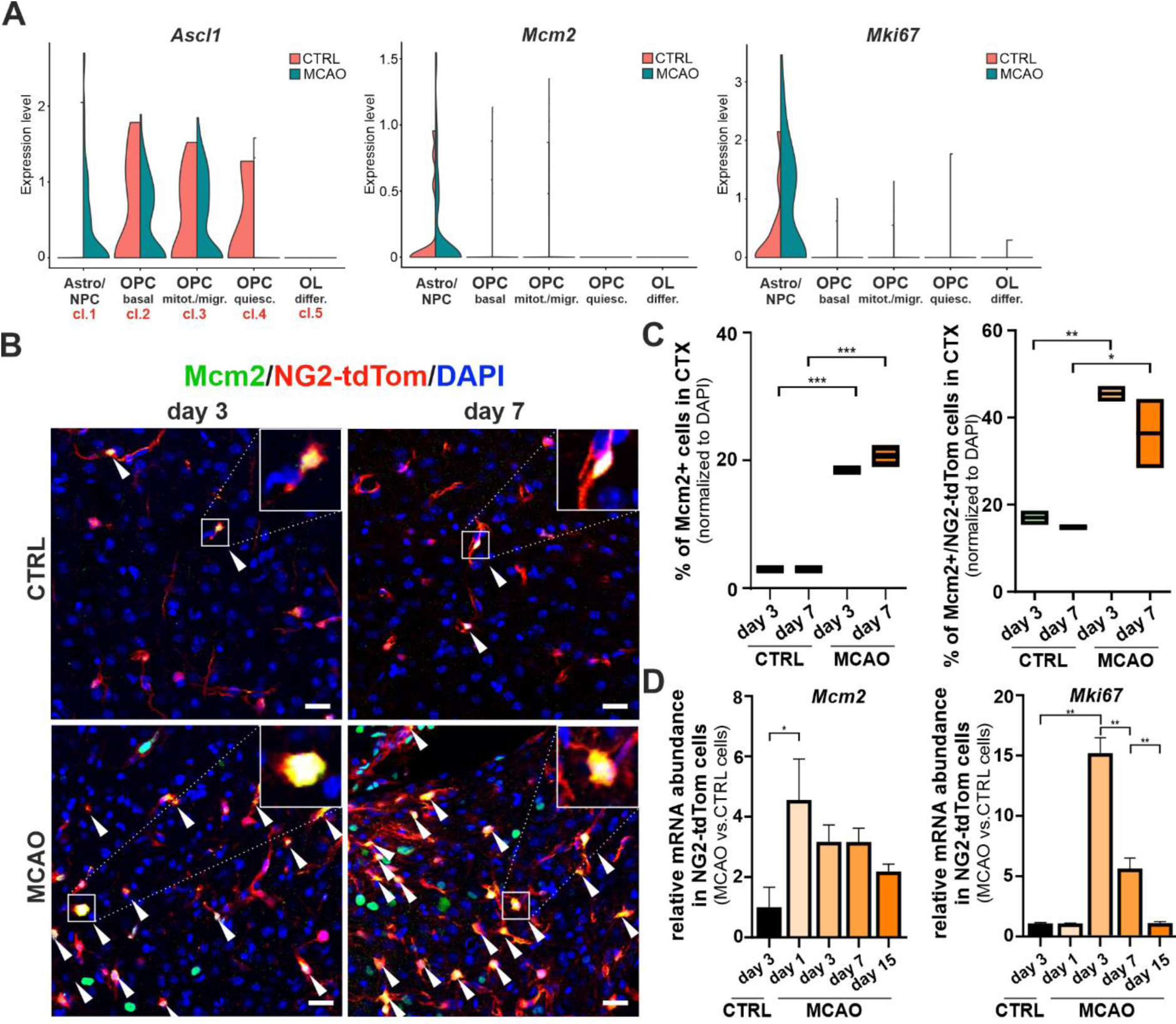
Examination of the expression of NSC and TAC genes in NG2-tdTom cells. (A) Violin plots showing expression of *Ascl1*, *Mcm2*, and *Mki67* genes in NG2-tdTom/Pdgfrα^+^ subpopulations identified in the cortex of the sham-operated (CTRL) or ischemic brain 3 days after MCAO. Cell cluster numbers, as indicated in the text and shown in Fig. 4B and Table 1, are listed in the upper left diagram. (B) Representative samples show Mcm2 and tdTomato positivity in the cortex around the ischemic wound and in the CTRL sample; colocalization of both fluorescence signals is indicated by white arrowheads. Enlarged images are shown in the insets. Scale bar indicates 20 μm. (C) Quantification of Mcm2^+^ (left) and Mcm2^+^/NG2-tdTom double-positive cells (right) in the ischemic and CTRL cortex. (D) Expression levels of *Mcm2* and *Mki67* genes in NG2-tdTom cells sorted at different days after MCAO; expression in the control sample (analyzed at day 3 after surgery) was arbitrarily set to 1. Error bars represent the SD of the technical triplicates. Statistical significance (C, D) was determined by one-way ANOVA test; * p < 0.05; ** p < 0.01; *** p < 0.001.

Mcm2 positivity with simultaneous expression of *Ascl1* and *GFAP* is the hallmark of active NSCs (Andersen et al. 2014, Sueda et al. 2019). Because the expression of both *Ascl1* and *Mcm2* genes is increased in the astro/NPC-like glia subpopulation after ischemic injury and, furthermore, *de novo* expression of *GFAP* is activated in these cells, we hypothesized that this subpopulation may have the potential to produce different cell types after ischemic injury. Therefore, we focused on the genes whose expression was increased in cells of the neurogenic lineage and simultaneously in NG2 glia. In contrast to CTRL tissue, we detected expression of the brain expressed X-linked (*Bex*), Purkinje cell protein 4 (*Pcp4*), small nucleolar RNA host gene 11 (*Snhg11*), and SRY-box transcription factor 11 (*Sox11*) genes in NG2-tdTom cells in brain tissue after MCAO (Fig. 8 and Suppl. Fig. S4B). The *Bex* gene expression has already been demonstrated in neurons in different parts of the brain (Koo, Saraswati, and Margolis 2005), and the function of these genes has been linked to neurodegenerative diseases (Fernandez, Diaz-Ceso, and Vilar 2015) and neuronal repair after injury (Khazaei et al. 2010). Our analysis showed that in CTRL brain tissue, mRNA expression of *Bex1* was found in basal and mitotic/migratory OPCs. The ischemic injury resulted in a slight increase in *Bex1* expression in OPCs and *de novo* expression in astro/NPC-like cells (Fig. 8B). We also observed the expression of *Pcp4*, which encodes a neurospecific calmodulin-binding protein. Pcp4 production has been associated with neuronal differentiation, neurotransmitter release, and blocking neuronal apoptosis (Harashima et al. 2011). Similar to *Bex1*, the expression of *Pcp4* in healthy tissues was restricted to OPCs, whereas in ischemic tissue it occurred in a subpopulation of astro/NPC-like NG2 glia (Fig. 8B). Two other genes analyzed were *Snhg11* and *Sox11*. The expression of Snhg11 was observed in different types of mature neurons in the adult mouse brain (Kiss et al. 2020, MacKay et al. 2019, Tan et al. 2013); Sox11 is required for proliferation and differentiation of NPCs during embryonic development and in the adult brain (Da Silva et al. 2021, Li et al. 2012, Wang et al. 2013). We detected *Snhg11* expression in OPCs in CTRL and ischemic tissues. In contrast, *Sox11* expression in CTRL mice was detectable in all subpopulations of NG2 glia; however, after MCAO, the expression of this gene disappeared from resting OPCs and differentiating OLs, but was increased in astro/NPC-like cells (Fig. 8B). The time course of changes in gene expression of *Bex1*, *Pcp4*, and S*ox11* after ischemic brain injury was similar. A significant increase in the expression of these genes was observed at days 3 and 7 after MCAO, followed by a significant decrease at day 15. In contrast, the expression of the *Snhg11* gene gradually increased from day 3 and reached the maximum at day 15 after MCAO (Fig. 8C). We detected the presence of Pcp4 at the protein level by immunohistochemical staining. Immunohistochemistry revealed large Pcp4-positive neurons in the ischemic cortex at day 3; however, these Pcp4-producing neurons were also present in healthy tissue. At later intervals after ischemia, the number of these neurons decreased. However, most NG2-tdTom cells near the ischemic injury showed positivity for Pcp4. Interestingly, at day 12 after MCAO, Pcp4-producing NG2-tdTom cells were localized at the edge of the ischemic scar (Fig. 8D).

**Figure 8.**
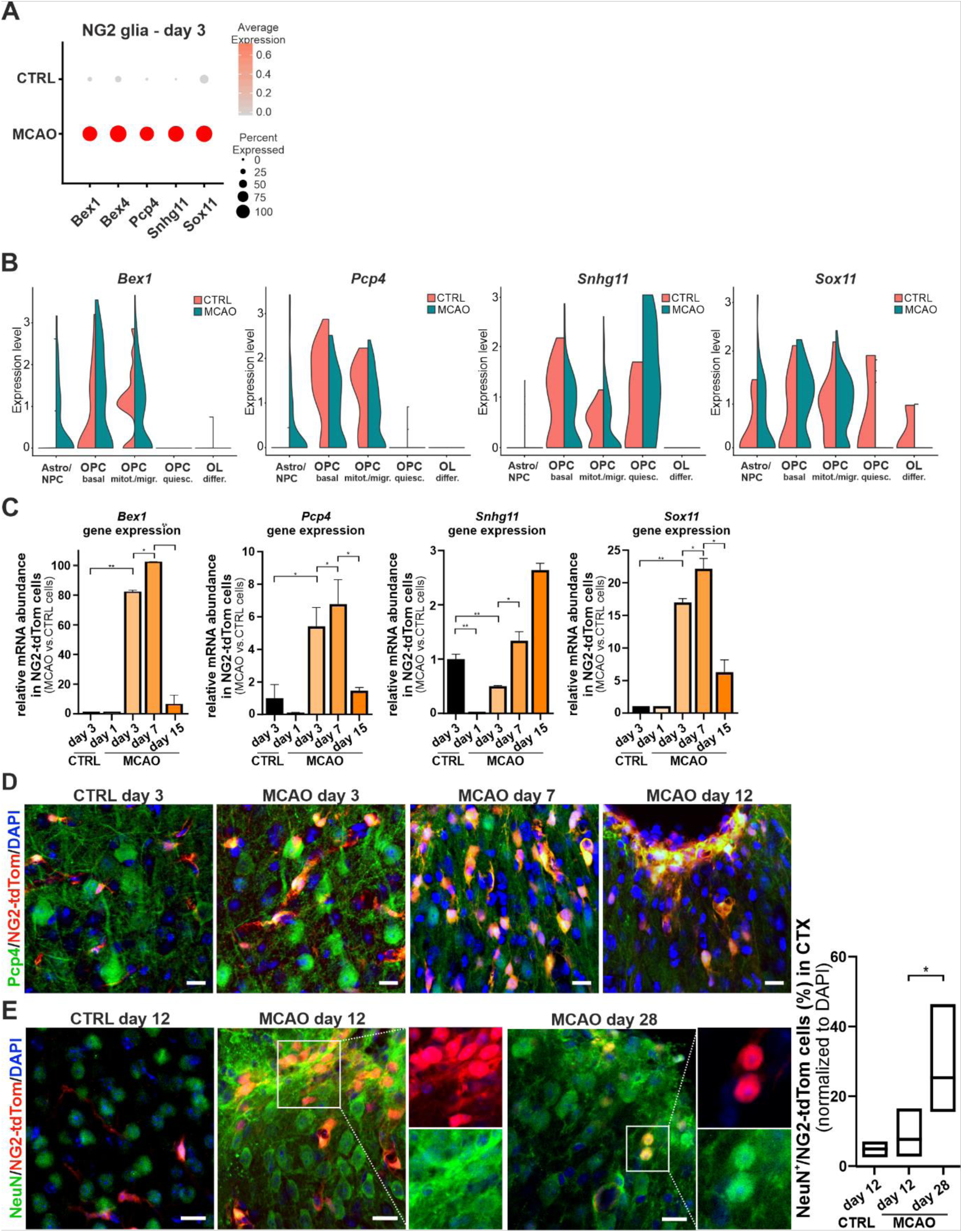
A subset of NG2-tdTom cells express markers involved in neuronal formation. (A) Dot plot shows expression of selected genes in NG2-tdTom/Pdgfrα^+^ cells in the cortex of the sham-operated (CTRL) or ischemic brain 3 days after MCAO. Color coding shows the increase in gene expression compared with the average gene expression of all NG2-tdTom/Pdgfrα^+^ cells; dot size corresponds to the percentage of cells expressing the selected gene in the given sample. (B) Violin plots showing expression of the indicated genes in the NG2-tdTom/Pdgfrα^+^ cell subpopulations. (C) Expression levels of the indicated genes in NG2-tdTom cells fluorescence-activated cell sorted at different days after MCAO. Expression in the CTRL sample obtained 3 days after surgery was arbitrarily set to 1. Error bars represent SD of technical triplicates. (D, E) Representative samples show colocalization of cells stained with Purkinje cell protein 4 Pcp4 (D) or neuron-specific nuclear protein NeuN (E) with endogenous fluorescence of NG2-tdTom cells near the ischemic wound at specific time points and in the corresponding sham-operated CTRL. Scale bars represent 10 μm in size. The graph on the right shows the colocalization of NeuN antibody staining with red fluorescent signal in NG2-tdTom cells. Statistical significance was determined using a one-way ANOVA test; *, p < 0.05; **, p < 0.01. Bex1/4, brain expressed X-linked gene 1/4; Pcp4, Purkinje cell protein 4; Snhg11, small nucleolar RNA host gene 11; Sox11, SRY-box transcription factor 11 (Sox11).

To answer the question whether NG2 glia in the ischemic cortex have the potential to differentiate into mature neurons, we used immunostaining for the NeuN protein. Although the function of the NeuN protein is not entirely clear, it is present in the nucleus and perinuclear cytoplasm of most post-mitotic neurons and has been used as a universal marker of neuronal differentiation for a decade (reviewed in (Gusel’nikova and Korzhevskiy 2015)). Whereas we detected a number of neurons, i.e., NeuN^+^ cells, and a small number of NG2-tdTom cells among them, in the healthy tissue, the distribution of neurons around the ischemic lesion in the cortex was markedly different. At day 12 after MCAO, we observed more NG2-tdTom cells among the neurons at the lesion edge, and their red fluorescence signal mostly did not overlap with the staining for NeuN. Finally, 28 days after MCAO, there were fewer NeuN^+^ neurons in the ischemic area, but we detected rare double-positive cells predominantly present at the margin of the lesion. The number of NG2-tdTom cells was rather low (∼20 cells per field of view), but approximately 23% of them were NeuN-positive (Fig. 8E). The red fluorescence detected in these NeuN^+^ cells at such a long interval after FCI induction suggests the ability of some NG2 glial cells to differentiate into mature neurons and compensate for the neuronal loss after ischemic injury.

In conclusion, we observed the expression of NPC- and neuron-specific genes in the astro/NPC-like subpopulation of NG2 glia after ischemia. Our results suggest that this subpopulation is involved in potential neuronal regeneration after brain injury.

## DISCUSSION

In this study, we investigated the possible role and differentiation potential of NG2 glia in ischemic brain injury. While the importance of NG2 glia in tissue regeneration after injury has been described many times, their plasticity in the adult brain remains an unresolved question. Using transgenic animals with tamoxifen-inducible production of fluorescent protein tdTomato in Cspg4/NG2-expressing cells and their progeny, we demonstrated an increase and localization of NG2 glia at the edge of the ischemic lesion in the cortex 3 and 7 days after MCAO. We also identified five subpopulations of NG2/Pdgfrα^+^ cells and described their membrane properties and characteristic gene expression. The previously identified astrocyte-like NG2 glia were present at low levels in the cortex of CTRL mice. After ischemic injury, these cells proliferate strongly and express a large number of genes typical of NPCs. Finally, we disclosed the potential of NG2 glial cells to differentiate into NeuN^+^ cells, albeit at a relatively low level.

The majority of the isolated NG2-tdTom cells, even in the ischemic samples, were multifunctional mural cells called perivascular cells. We explain this by the fact that we sampled a larger region of the cortex of the ischemic hemisphere and not just the area around the lesion, which is rich in NG2 glia. When we compared the changes in the abundance of each cell type, we found that the total number of perivascular cells remained almost the same after ischemic stroke, compared with healthy tissue. This observation is interesting because perivascular cells have been described to proliferate and migrate in response to brain injury (see (Cai et al. 2017)). However, it has been shown that between days 3 and 11 after spinal cord injury, only about one-third of proliferating perivascular cells carry the NG2 marker and that the number of proliferating NG2 glia exceeds the number of proliferating NG2^+^ perivascular cells by a factor of 20 (Hesp et al. 2018). Perivascular cells appear to serve multiple functions in lesion healing, and their more detailed study certainly deserves attention, although this is beyond the scope of this study.

In the OL population, a slight decrease in their abundance after ischemic stroke could be explained by a balance between cell death of OLs after ischemic injury because of their high sensitivity to oxidative stress and excitotoxicity during cerebral ischemia (discussed in (Belov Kirdajova et al. 2020)) and the increased formation of new OLs observed in animals after MCAO. Although previous studies reported an increasing number of newly formed OLs within one to two weeks after stroke (Gregersen et al. 2001, Mabuchi et al. 2000), we observed a decrease in OLs derived from NG2-tdTom OPCs. Histological examination of the ischemic tissue confirmed that mature OLs formed rapidly at the injury site 7 days after MCAO; however, only a portion of these OLs showed red fluorescence at the same time. Therefore, it is evident that these OLs arose from OPCs with an inactive reporter allele, as a result of incomplete recombination of the reporter allele, or by the emergence of these OPCs from a source other than NG2-positive cells such as NSCs in the SVZ or mesenchymal stem cells (Dang et al. 2019).

The number of NG2 glia, which was the third most abundant population, increased 3 and 7 days after the onset of FCI but, surprisingly, was reduced in both ischemic and sham-operated animals at day 12. NG2 glia in the cortex continuously differentiate in response to brain injury, primarily into new OLs and to a lesser extent into other cell types (Kang et al. 2010, Rivers et al. 2008, Young et al. 2013). However, it has been described previously that NG2 glia maintain a stable population through homeostatic control of cell density, and loss of NG2 glia is continuously compensated by their proliferation. After injury, this constant turnover is further enhanced and supplemented by migration from other brain regions (Hughes et al. 2013). Therefore, the decreasing number of NG2-tdTom glia over a prolonged period after ischemia may reflect the transition from the acute response to injury to the phase of healing and partial recovery. Nonetheless, the rapid increase in the number of NG2 glia in the cortex shortly after MCAO and their accumulation at the edge of the lesion is consistent with the current knowledge of NG2 glia proliferation and their contribution to glial scar formation (reviewed in (Adams and Gallo 2018)).

Single-cell RNA-seq analysis of all isolated NG2-tdTom cells revealed a division into five distinct clusters of NG2 glia. Consistently, heterogeneity of adult brain NG2 glia has been reported recently (Beiter et al. 2022). In the subsequent Discussion section, we attempt to assign the electrophysiological parameters shown in Fig. 5 to the individual cell clusters. Because the measurements were performed on live cells labeled with endogenous red fluorescence, we could not take advantage of simultaneous labeling with an antibody that recognizes a specific marker of each cluster. Genetic labeling of different cell types would be possible but difficult because it would require crossing (or making *de novo*) specific genetically modified mouse strains. For this reason, we assigned these parameters to each cell group based on the available literature and our previous experience. However, we are aware that this assignment may not be entirely accurate, and possibly not all cells in a given cluster have the same electrophysiological properties.

The first cell cluster was identified as undifferentiated NG2 glia with a “complex” electrophysiological profile, i.e., with a large permeability to K^+^ ions, both inwardly and outwardly rectified K^+^ currents, and currents with a voltage-dependent character (Chittajallu, Aguirre, and Gallo 2004, Kriska et al. 2021). The passive membrane properties attributed to NG2 glia by Chittajallu and colleagues are not consistent with our results, i.e., the cells studied were more depolarized and had a lower C_M_. However, our measurements were performed on the adult cerebral cortex damaged by ischemia, whereas Chittajallu and colleagues studied healthy juvenile animals. Alterations in the membrane properties of NG2 glia in the ischemic brain have been described previously and included a depolarized V_M_ and a reduced C_M_ in rats after global ischemia (Pivonkova et al. 2010). When comparing passive membrane properties in healthy and ischemic tissues, we observed similar trends. We believe that this group of cells represents a subpopulation of basal OPCs, as they express many genes characteristic of these cells in different parts of the brain (Mizrak et al. 2019, Zhang et al. 2014).

The cells of the second cluster expressed most of the OPC-specific markers, like the other subpopulations, but differed in the expression of genes related to mitosis and cell migration. The expression of *Dcx* has been proposed as a marker of migrating OPCs (Boulanger and Messier 2017a) and was recently observed in SVZ-derived cells with similar membrane properties, including the absence of K_IR_, a high IR, and a depolarized V_M_ (Kriska et al. 2021). K_IR_ currents in NG2 glia are predominantly mediated by Kir4.1, which is highly active in the majority of NG2 glia and is thought to play a fundamental role in their development, survival, and differentiation (reviewed in (Larson, Zhang, and Bergles 2016)). Accordingly, the expression of potassium inwardly rectifying channel subfamily J member 10 (*Kcnj10*), which encodes Kir4.1, was decreased two-fold in this cluster compared to basal NG2 glia (not shown). Decreased K_IR_ currents have already been described in postnatal NG2 white matter glia (Chittajallu, Aguirre, and Gallo 2004), whose migration to the cortex may be triggered by injury. However, the absence of K_IR_ currents has not been observed in adult white matter NG2 glia (Larson et al. 2018), suggesting that this feature is restricted to developing NG2 glia. On the other hand, the absence of K_IR_ currents could also be related to cell cycle progression and proliferation status. A reduction in K_IR_ currents has been observed in proliferating reactive astrocytes after spinal cord injury (MacFarlane and Sontheimer 1997), cortical stab wound (Anderova et al. 2004) and freeze lesion (Bordey et al. 2001); and furthermore, cell cycle progression was impaired by K_IR_ manipulation at several checkpoints (MacFarlane and Sontheimer 2000). A similar phenomenon was also observed in retinal gliosis (Ulbricht et al. 2008). These findings and the absence of expression of genes characteristic of G1/S phase, such as *Mcm2* (Stoeber et al. 2001) or *Rrm2* (Zhang et al. 2020), led us to hypothesize that these may be cells that have recently undergone cell division and are migrating to a target site, i.e., (post-)mitotic/migratory OPCs. In the brain, a decrease in K_IR_ currents in NG2 glia was observed after injury following both global (Pivonkova et al. 2010) and focal cerebral ischemia (Song et al. 2018). Consistent with this, we recorded an enrichment of the mitotic/migratory OPC population at the expense of “complex” basal NG2 glia in the sample 3 days after MCAO compared to healthy tissue. We conclude that this change may reflect increased migration of NG2 glia to the site of injury.

The third cluster of cells was identified only by differential gene expression. We were unable to distinguish these cells on the basis of their electrophysiological properties, either because their V_M_ is not significantly different from that of basal or mitotic/migratory OPCs, or their abundance in ischemic tissue is low. This could also be the reason why the two previous publications discussing NG2 glia subtypes did not distinguish this minority group of OPCs (Kirdajova et al. 2021, Valny et al. 2018). Another reason could be their possibly different morphology, which would exclude them from the patch-clamp analysis as non-NG2 glial cells. Considering the overall lower expression of most marker genes, we conclude that these cells may be quiescent and belong to a stable, non-dividing population that does not change during physiological aging of the brain (Psachoulia et al. 2009), but is activated after injury. Moreover, the genes expressed in this subset of NG2 glia have been assigned by GSEA to the compartment of the axon initial segment. Therefore, it is possible that these cells serve a specific function, e.g., in communication of NG2 glia with neurons (discussed in (Yang, Xiong, and Yao 2013)), which would correlate with their loss in the cortex after FCI.

A subpopulation of NG2 glia destined for the OL lineage was localized to the fourth cell cluster based on the OL-differentiation-specific genes and GSEA. In the patch-clamp analysis, these cells showed time- and voltage-independent decaying K^+^ currents, similar to differentiated OLs (Honsa et al. 2012). The passive membrane properties of these committed OL progenitor cells differed slightly from those described by Honsa and colleagues, as they exhibited a higher IR and lower K_IR_ and K_DR_. This could be due to a different degree of differentiation into OLs, but the membrane properties of the committed OL progenitor cells were obviously different from astrocytes or oligodendrocyte precursors and were more similar to the membrane properties of mature oligodendrocytes described previously (Chvatal et al. 1999). Of note, the level of K_IR_ currents further increases again during oligodendrocyte maturation, as Kir4.1 is essential for the myelination process (Hong et al. 2023). Consistently, we observed increasing expression of *Kcnj10* in mature OL clusters. As previously described (Kirdajova et al. 2021), the number of committed OL progenitor cells decreases in the early response to FCI. This may be due to the fact that NG2 glia that differentiate into OLs are more sensitive to ischemic injury than mature OLs.

According to the GSEA, the fifth cell cluster corresponded to the NPCs of the prefrontal cortex. Cells in this cluster clearly expressed the astrocytic markers *Cldn10* and *GFAP*, markers of active neural progenitors *Egfr*, nestin, and vimentin, and a number of cell cycle-related genes such as *Mki67*, *Mcm2*, *Spc24*, and *Ube2c*. When compared with genes specific to NSCs and TACs (Kriska et al. 2021), we also found a considerable overlap. In terms of electrophysiological properties, these cells were characterized by passive time- and voltage-independent non-decaying K^+^ currents, previously described in astrocytes (Anderova et al. 2004, Kriska et al. 2021, Zhou, Schools, and Kimelberg 2006), which is why we termed this subpopulation “astro/NPC-like”. Cells with similar passive membrane properties were previously observed in NG2-derived cells; importantly, these cells were characterized by *GFAP* gene expression and were therefore considered “astrocyte-like” (Honsa et al. 2012, Kirdajova et al. 2021). The astrocyte-like subpopulation of NG2 glia was previously described as transient only in ischemic tissue. A well-known response to tissue damage in the CNS, reactive astrogliosis, is a process in which the number of astrocytes at the ischemic site increases and their morphology changes due to remodeling of intermediate filaments composed mainly of GFAP, vimentin, and nestin (reviewed in (Hol and Pekny 2015)). The proliferation of GFAP^+^ cells and the increase in their number in the ischemic cortex after MCAO is consistent with this mechanism. Importantly, in the ischemic tissue we found NG2-tdTom cells that were positive for both GFAP and vimentin. Simultaneous expression of these genes together with nestin was detected only in the astro/NPC-like subpopulation of NG2 glia, although vimentin was also expressed in OPCs. Furthermore, the gene expression of astrocyte-like NG2 cells reported by Kirdajova and colleagues (Kirdajova et al. 2021) is consistent with that of astro/NPC-like cells, and we, therefore, speculate that they are the same subpopulation of NG2 glia. Moreover, the electrophysiological properties of the astro/NPC-like cells resembled those observed in GFAP^+^ nestin^+^ cells in the hippocampus of adult mice. Interestingly, these cells were not positive for astrocytic marker S100β (Filippov et al. 2003). Therefore, they might rather represent mitotically active NPCs sharing some properties with astrocytes (Seri et al. 2001). Accordingly, GFAP has been repeatedly described to be present in the subpopulations of adult astrocytes that give rise to neurons, in addition to reactive astrocytes (Casper and McCarthy 2006, Doetsch, Garcia-Verdugo, and Alvarez-Buylla 1997, Garcia et al. 2004). Our previous scRNA-seq analysis of SVZ+S cells (Kriska et al. 2021) confirmed the presence of GFAP in astro-to-NSC transition cells in this region. However, cortical astrocytes in the healthy adult brain show less GFAP positivity than in neurogenic regions such as the hippocampus (Zhang et al. 2019). Therefore, it is likely that the presence of GFAP^+^ NPCs in the cortex is related to brain injury. Importantly, gene expression analysis of all (*in silico* unsorted) cells allowed us to also assign healthy cerebral cortex cells to this subpopulation, which, unlike those in the ischemic tissue, do not express GFAP (our observation). On the basis of the gene signature of the progenitor cells and the fact that the expression of cell cycle-specific genes is restricted to these cells, we propose that the astro/NPC-like subpopulation of NG2 glia serves as a source of NG2 glia replenishment and is thus critical for NG2 glia self-renewal. This would be consistent with the increased numbers of these cells in the ischemic cortex. At the same time, Mcm2 expression was relatively low in the healthy cortex, but among NG2-tdTom cells, nearly 20% expressed this cell proliferation marker, supporting the notion that NG2 glia represent the major proliferating cell type in the adult healthy brain (Dawson et al. 2003). Although the number of Mcm2^+^ cells in the ischemic cortex increased slightly over time, the number of Mcm2^+^ cells among NG2-tdTom cells was lower at day 7 after FCI than at day 3. Therefore, NG2 glia appear to be responsible for the initial proliferative response to injury. Strikingly, in a recent paper, Delgado and colleagues demonstrated the existence of NG2^+^ Pdgfrα^+^ cells, some of which were GFAP-positive and many of which carried both Mcm2 and Ki67 markers. These cells were located in the ventricle on the inner side of the SVZ, whereas no OL differentiation was observed in this region. After focal demyelination, the number of these intraventricular cells increased in response to brain injury (Delgado et al. 2021), which is completely consistent with what we observed in the ischemic cortex. Therefore, it is likely that these astro/NPC-like cells derived from NG2 glia are not restricted to the cortex, or perhaps these cells migrate to (any) injured region. The same study also showed that many of the detected intraventricular NG2 glia were positive for Ascl1. Interestingly, we observed that *Ascl1* expression occurred in astro/NPC-like cells (Fig. 7A). The *Ascl1* gene (also known as *Mash1*) encodes a transcription factor that has been previously linked to neurogenesis (see (Vasconcelos and Castro 2014)).

The gene has been successfully used to reprogram non-neuronal cells into neurons *in vitro* (Berninger et al. 2007, Vierbuchen et al. 2010). *In vivo*, however, the situation seems to be more complicated, as some studies report that Ascl1 alone is not sufficient to induce redifferentiation (Grande et al. 2013), while others suggest the opposite (Liu et al. 2015). Efforts have been made to reprogram NG2 glia *in vivo*, and Ascl1 was one of the three factors used to induce differentiation of NG2 glia into NeuN^+^ and MAP2^+^ cells. Cortical NG2 glia were also successfully reprogrammed into neurons by increased *Sox2* and *Ascl1* expression, but this conversion was only successful in tissues after injury (Heinrich et al. 2014). Increased *Ascl1* expression was observed in NG2 glia after spinal cord injury; however, this increased expression did not induce neurogenesis unless *Sox2* was also highly expressed. In fact, inducible expression of *Sox2* further increased *Ascl1* expression (Tai et al. 2021), suggesting that endogenous Sox2 and Ascl1 levels in NG2 glia are not sufficient to induce neurogenesis. Consistent with this, *Ascl1* expression in NG2 glia was shown to be important for their proliferation and maintenance; overexpression of *Ascl1* decreased OL differentiation and, in turn, induced neuronal differentiation (Sueda et al. 2019). We detected *Sox2* mRNA in all subtypes of NG2 glia, whose abundance gradually decreased with OL maturation independent of ischemia. This suggests that Sox2 is involved in maintaining the progenitor status of NG2 glia, but is not a direct or sole regulator of *Ascl1* expression and, unlike Ascl1, is not responsive to ischemic tissue damage.

Other genes expressed in astro/NPC-like cells after MCAO were *Bex1* and *Pcp4*. Bex genes were previously associated with the regulation of apoptosis in oligodendrocytes (Kimura et al. 2001); later studies reported their role in cell cycle, neuroprotection (Vilar et al. 2006) and axonal regeneration (Khazaei et al. 2010). In addition, increased expression of this gene has been observed in retinal ganglion cells following optic nerve stroke (Bernstein et al. 2006). The function of Pcp4, also known as peptide 19 (Pep19), has been associated with neuronal survival (Kanazawa et al. 2008) and neurotransmitter release (Harashima et al. 2011). Interestingly, knocking down *Bex1* accelerated neuronal differentiation *in vitro* (Vilar et al. 2006); therefore, the decrease in its expression in NG2 glia over a prolonged period after FCI may be related to injury-induced neurogenesis. On the other hand, neuronal differentiation was observed when *Pcp4* was overexpressed during embryogenesis (Mouton-Liger et al. 2011). In contrast to the decrease in *Pcp4* mRNA levels, histological examination of the tissue 12 days after MCAO revealed that almost all NG2-tdTom cells at the edge of the lesion were positive for Pcp4 protein, suggesting a role for Pcp4 in NG2 glia directly at the site of injury. It should be mentioned that we observed a decrease in the number of NG2 glia among NG2-tdTom cells as early as 12 days after MCAO; therefore, the decrease in *Pcp4* (and *Bex1*) gene expression at a later time point after MCAO could be due to the increased proportion of perivascular cells in the isolated sample. Interestingly, Pcp4 might upregulate the expression of *Ascl1* (Kitazono et al. 2020). Nevertheless, to our knowledge, the function of Pcp4 in NG2 glia has not been elucidated. Pcp4 is involved in calmodulin-dependent signaling by direct binding of calmodulin (Putkey et al. 2008); Ca^2+^-dependent calmodulin binding by Bex1 has been demonstrated in skeletal muscle (Koo et al. 2007). Ca^2+^/calmodulin signaling plays an essential role in neuroprotection against cerebrovascular injury, as calmodulin-dependent kinase knockout mice exhibit more severe brain damage and behavioral deficits after ischemic stroke (Liu, McCullough, and Li 2014, McCullough et al. 2013). It is therefore possible that the increased expression of both genes shortly after the onset of FCI is related to this neuroprotective function. Similarly, the increased expression of *Snhg11* in NG2-tdTom/Pdgfrα^+^ cells after MCAO may be due to tissue protection, as this gene plays a role in regulating damage caused by oxygen-glucose deprivation/reperfusion in an *in vitro* model of ischemia (Chen, Fan, and Wu 2021). Another gene that is upregulated in ischemic tissue shortly after FCI, *Sox11*, has been reported to be essential for initiating neuronal expression in undifferentiated NPCs (Bergsland et al. 2006). Moreover, its role in neuroregeneration after injury has already been described in several studies (Guo et al. 2014, Jankowski et al. 2009, Jing et al. 2012, Struebing et al. 2017). Finally, the observed decrease in *Sox11* expression two weeks after MCAO might be related to the fact that its reduced expression is critical for the completion of OL differentiation (Swiss et al. 2011).

The expression of genes associated with neurogenesis, neuroprotection, and neuroregeneration and of factors associated with active NSCs suggests a possible neurogenic potential of the astro/NPC-like NG2 glia subpopulation in the ischemic cortex. Although we found NG2-tdTom cells that were positive for the neuronal marker NeuN 28 days after MCAO, we assume that gene expression *per se* does not represent evidence that functional neurons can arise from the progeny of NG2-positive cells. Given the relatively small number of NG2-tdTom/NeuN-positive cells, there is a possibility of genetic labeling of neurons facilitated by Cre recombinase transfer from NG2-positive cells to neurons via exosomes, turning on the expression of the tdTomato reporter protein in the Cspg4/NG2-negative cells (Fruhbeis et al. 2013). Furthermore, due to their low abundance, we could not assess whether these NeuN^+^ cells form functional synapses and integrate into neuronal circuits. Nonetheless, our results demonstrate “neurogenic” gene expression along with the presence of NeuN positivity in NG2-tdTom cells after ischemic injury, supporting previous studies suggesting that NG2 glia may differentiate into neurons under certain/specific conditions (Boulanger and Messier 2017b, Rivers et al. 2008, Robins, Trudel, et al. 2013). In the future, it would be advantageous to find a specific marker for the astro/NPC-like NG2 glia subpopulation in order to monitor its behavior and progeny in the brain after injury.

In conclusion, NG2 glia perform multiple functions in the response to brain injury and in tissue regeneration around the ischemic lesion. Our data suggest that astro/NPC-like NG2 glia represent a progenitor pool for OPCs and that they may acquire neurogenic potential after an ischemic insult. However, the capacity of NG2 glia to form new neurons is relatively low, and we might speculate that the expression of key factors inducing neurogenesis is insufficient or that neuronal differentiation is suppressed by the local microenvironment of the cerebral cortex. However, the neurogenic capacity of astro/NPC-like NG2 glia suggests a dormant potential of these cells. Discovering ways to harness this intrinsic potential may provide new approaches to promote neuronal tissue regeneration and thus improve recovery after ischemic CNS injury.

## ACKNOWLEDGEMENTS

We thank Sarka Takacova for critical reading of the manuscript, and Marketa Hemerova Helena Pavlikova and Sarka Kocourkova for their technical assistance. This research was funded by the Czech Science Foundation, grant no. 21-24674S, and by the Czech Academy of Sciences (Strategy AV21, grant no. VP29). JKr was supported by project L200392251 from the Czech Academy of Sciences; TK was in part supported by the Grant Agency of Charles University (grant no. 330119). MK and JKu were in part supported by the Ministry of Education, Youth and Sports (ELIXIR CZ - LM2018131). The Light Microscopy Core Facility, IMG CAS, Prague, Czech Republic, was supported by the Ministry of Education, Youth and Sports (LM2018129, CZ.02.1.01/0.0/0.0/18_046/0016045) and RVO – 68378050-KAV-NPUI.

## AUTHOR CONTRIBUTIONS

MA and VK designed the study concept and supervised the project. MA, VK, TK, and LJ coordinated the experiments. TK, JKr, ZH, and JT were responsible for the tissue isolation and processing; TK, SCG, and DK performed MCAO operations. JKu and MK carried out bioinformatics processing; LJ and LB facilitated analysis of the gene expression data from scRNA-seq and RT-qPCR experiments. TK and JKr performed the electrophysiological study. TK and LJ acquired and analyzed the data from the immunochemical experiments. LJ and KG supervised the breeding of the transgenic mouse strains. TK and LJ wrote the manuscript. JKr, VK, and MA revised the manuscript. All authors read and approved the final version of the manuscript.

## REFERENCES

Adams, K. L., and V. Gallo. 2018. “The diversity and disparity of the glial scar.” Nat Neurosci 21 (1):9–15. doi: 10.1038/s41593-017-0033-9.

Akay, L. A., A. H. Effenberger, and L. H. Tsai. 2021. “Cell of all trades: oligodendrocyte precursor cells in synaptic, vascular, and immune function.” Genes Dev 35 (3-4):180–198. doi: 10.1101/gad.344218.120.

Anderova, M., T. Antonova, D. Petrik, H. Neprasova, A. Chvatal, and E. Sykova. 2004. “Voltage-dependent potassium currents in hypertrophied rat astrocytes after a cortical stab wound.” Glia 48 (4):311–26. doi: 10.1002/glia.20076.

Andersen, J., N. Urban, A. Achimastou, A. Ito, M. Simic, K. Ullom, B. Martynoga, M. Lebel, C. Goritz, J. Frisen, M. Nakafuku, and F. Guillemot. 2014. “A transcriptional mechanism integrating inputs from extracellular signals to activate hippocampal stem cells.” Neuron 83 (5):1085–97. doi: 10.1016/j.neuron.2014.08.004.

Androvic, P., D. Kirdajova, J. Tureckova, D. Zucha, E. Rohlova, P. Abaffy, J. Kriska, M. Valny, M. Anderova, M. Kubista, and L. Valihrach. 2020. “Decoding the Transcriptional Response to Ischemic Stroke in Young and Aged Mouse Brain.” Cell Rep 31 (11):107777. doi: 10.1016/j.celrep.2020.107777.

Ashburner, M., C. A. Ball, J. A. Blake, D. Botstein, H. Butler, J. M. Cherry, A. P. Davis, K. Dolinski, S. S. Dwight, J. T. Eppig, M. A. Harris, D. P. Hill, L. Issel-Tarver, A. Kasarskis, S. Lewis, J. C. Matese, J. E. Richardson, M. Ringwald, G. M. Rubin, and G. Sherlock. 2000. “Gene ontology: tool for the unification of biology. The Gene Ontology Consortium.” Nat Genet 25 (1):25–9. doi: 10.1038/75556.

Becht, E., L. McInnes, J. Healy, C. A. Dutertre, I. W. H. Kwok, L. G. Ng, F. Ginhoux, and E. W. Newell. 2018. “Dimensionality reduction for visualizing single-cell data using UMAP.” Nat Biotechnol. doi: 10.1038/nbt.4314.

Beiter, R. M., C. Rivet-Noor, A. R. Merchak, R. Bai, D. M. Johanson, E. Slogar, K. Sol-Church, C. C. Overall, and A. Gaultier. 2022. “Evidence for oligodendrocyte progenitor cell heterogeneity in the adult mouse brain.” Sci Rep 12 (1):12921. doi: 10.1038/s41598-022-17081-7.

Belov Kirdajova, D., J. Kriska, J. Tureckova, and M. Anderova. 2020. “Ischemia-Triggered Glutamate Excitotoxicity From the Perspective of Glial Cells.” Front Cell Neurosci 14:51. doi: 10.3389/fncel.2020.00051.

Bergles, D. E., J. D. Roberts, P. Somogyi, and C. E. Jahr. 2000. “Glutamatergic synapses on oligodendrocyte precursor cells in the hippocampus.” Nature 405 (6783):187–91. doi: 10.1038/35012083.

Bergsland, M., M. Werme, M. Malewicz, T. Perlmann, and J. Muhr. 2006. “The establishment of neuronal properties is controlled by Sox4 and Sox11.” Genes Dev 20 (24):3475–86. doi: 10.1101/gad.403406.

Bernal, G. M., and D. A. Peterson. 2011. “Phenotypic and gene expression modification with normal brain aging in GFAP-positive astrocytes and neural stem cells.” Aging Cell 10 (3):466–82. doi: 10.1111/j.1474-9726.2011.00694.x.

Berninger, B., M. R. Costa, U. Koch, T. Schroeder, B. Sutor, B. Grothe, and M. Gotz. 2007. “Functional properties of neurons derived from in vitro reprogrammed postnatal astroglia.” J Neurosci 27 (32):8654–64. doi: 10.1523/JNEUROSCI.1615-07.2007.

Bernstein, S. L., J. H. Koo, B. J. Slater, Y. Guo, and F. L. Margolis. 2006. “Analysis of optic nerve stroke by retinal Bex expression.” Molecular Vision 12 (16):147–155.

Biname, F. 2014. “Transduction of Extracellular Cues into Cell Polarity: the Role of the Transmembrane Proteoglycan NG2.” Molecular Neurobiology 50 (2):482–493. doi: 10.1007/s12035-013-8610-8.

Binder, J. X., S. Pletscher-Frankild, K. Tsafou, C. Stolte, S. I. O’Donoghue, R. Schneider, and L. J. Jensen. 2014. “COMPARTMENTS: unification and visualization of protein subcellular localization evidence.” Database (Oxford) 2014:bau012. doi: 10.1093/database/bau012.

Boda, E., F. Vigano, P. Rosa, M. Fumagalli, V. Labat-Gest, F. Tempia, M. P. Abbracchio, L. Dimou, and A. Buffo. 2011. “The GPR17 receptor in NG2 expressing cells: focus on in vivo cell maturation and participation in acute trauma and chronic damage.” Glia 59 (12):1958–73. doi: 10.1002/glia.21237.

Bonfanti, E., P. Gelosa, M. Fumagalli, L. Dimou, F. Vigano, E. Tremoli, M. Cimino, L. Sironi, and M. P. Abbracchio. 2017. “The role of oligodendrocyte precursor cells expressing the GPR17 receptor in brain remodeling after stroke.” Cell Death Dis 8 (6):e2871. doi: 10.1038/cddis.2017.256.

Bordey, A., S. A. Lyons, J. J. Hablitz, and H. Sontheimer. 2001. “Electrophysiological characteristics of reactive astrocytes in experimental cortical dysplasia.” J Neurophysiol 85 (4):1719–31. doi: 10.1152/jn.2001.85.4.1719.

Boulanger, J. J., and C. Messier. 2017a. “Doublecortin in Oligodendrocyte Precursor Cells in the Adult Mouse Brain.” Front Neurosci 11:143. doi: 10.3389/fnins.2017.00143.

Boulanger, J. J., and C. Messier. 2017b. “Unbiased stereological analysis of the fate of oligodendrocyte progenitor cells in the adult mouse brain and effect of reference memory training.” Behav Brain Res 329:127–139. doi: 10.1016/j.bbr.2017.04.027.

Cai, W., H. Liu, J. Zhao, L. Y. Chen, J. Chen, Z. Lu, and X. Hu. 2017. “Pericytes in Brain Injury and Repair After Ischemic Stroke.” Transl Stroke Res 8 (2):107–121. doi: 10.1007/s12975-016-0504-4.

Casper, K. B., and K. D. McCarthy. 2006. “GFAP-positive progenitor cells produce neurons and oligodendrocytes throughout the CNS.” Mol Cell Neurosci 31 (4):676–84. doi: 10.1016/j.mcn.2005.12.006.

Chen, E. Y., C. M. Tan, Y. Kou, Q. Duan, Z. Wang, G. V. Meirelles, N. R. Clark, and A. Ma’ayan. 2013. “Enrichr: interactive and collaborative HTML5 gene list enrichment analysis tool.” BMC Bioinformatics 14:128. doi: 10.1186/1471-2105-14-128.

Chen, Y., Z. Fan, and Q. Wu. 2021. “Dexmedetomidine improves oxygen-glucose deprivation/reoxygenation (OGD/R) -induced neurological injury through regulating SNHG11/miR-324-3p/VEGFA axis.” Bioengineered 12 (1):4794–4804. doi: 10.1080/21655979.2021.1957071.

Chittajallu, R., A. Aguirre, and V. Gallo. 2004. “NG2-positive cells in the mouse white and grey matter display distinct physiological properties.” J Physiol 561 (Pt 1):109–22. doi: 10.1113/jphysiol.2004.074252.

Chvatal, A., M. Anderova, D. Ziak, and E. Sykova. 1999. “Glial depolarization evokes a larger potassium accumulation around oligodendrocytes than around astrocytes in gray matter of rat spinal cord slices.” J Neurosci Res 56 (5):493–505. doi: 10.1002/(SICI)1097-4547(19990601)56:5<493::AID-JNR5>3.0.CO;2-O.

Clarke, L. E., K. M. Young, N. B. Hamilton, H. Li, W. D. Richardson, and D. Attwell. 2012. “Properties and fate of oligodendrocyte progenitor cells in the corpus callosum, motor cortex, and piriform cortex of the mouse.” J Neurosci 32 (24):8173–85. doi: 10.1523/JNEUROSCI.0928-12.2012.

Cochard, L. M., L. C. Levros, Jr., S. E. Joppe, F. Pratesi, A. Aumont, and K. J. L. Fernandes. 2021. “Manipulation of EGFR-Induced Signaling for the Recruitment of Quiescent Neural Stem Cells in the Adult Mouse Forebrain.” Front Neurosci 15:621076. doi: 10.3389/fnins.2021.621076.

Colak, G., A. J. Filiano, and G. V. Johnson. 2011. “The application of permanent middle cerebral artery ligation in the mouse.” J Vis Exp (53). doi: 10.3791/3039.

Da Silva, F., K. Zhang, A. Pinson, E. Fatti, M. Wilsch-Brauninger, J. Herbst, V. Vidal, A. Schedl, W. B. Huttner, and C. Niehrs. 2021. “Mitotic WNT signalling orchestrates neurogenesis in the developing neocortex.” EMBO J 40 (19):e108041. doi: 10.15252/embj.2021108041.

Dang, T. C., Y. Ishii, V. Nguyen, S. Yamamoto, T. Hamashima, N. Okuno, Q. L. Nguyen, Y. Sang, N. Ohkawa, Y. Saitoh, M. Shehata, N. Takakura, T. Fujimori, K. Inokuchi, H. Mori, J. Andrae, C. Betsholtz, and M. Sasahara. 2019. “Powerful Homeostatic Control of Oligodendroglial Lineage by PDGFRalpha in Adult Brain.” Cell Rep 27 (4):1073–1089 e5. doi: 10.1016/j.celrep.2019.03.084.

Dawson, M. R., A. Polito, J. M. Levine, and R. Reynolds. 2003. “NG2-expressing glial progenitor cells: an abundant and widespread population of cycling cells in the adult rat CNS.” Mol Cell Neurosci 24 (2):476–88. doi: 10.1016/s1044-7431(03)00210-0.

Dayer, A. G., K. M. Cleaver, T. Abouantoun, and H. A. Cameron. 2005. “New GABAergic interneurons in the adult neocortex and striatum are generated from different precursors.” J Cell Biol 168 (3):415–27. doi: 10.1083/jcb.200407053.

de Pablo, Y., M. Nilsson, M. Pekna, and M. Pekny. 2013. “Intermediate filaments are important for astrocyte response to oxidative stress induced by oxygen-glucose deprivation and reperfusion.” Histochem Cell Biol 140 (1):81–91. doi: 10.1007/s00418-013-1110-0.

Delgado, A. C., A. R. Maldonado-Soto, V. Silva-Vargas, D. Mizrak, T. von Kanel, K. R. Tan, A. Paul, A. Madar, H. Cuervo, J. Kitajewski, C. S. Lin, and F. Doetsch. 2021. “Release of stem cells from quiescence reveals gliogenic domains in the adult mouse brain.” Science 372 (6547):1205–1209. doi: 10.1126/science.abg8467.

Diers-Fenger, M., F. Kirchhoff, H. Kettenmann, J. M. Levine, and J. Trotter. 2001. “AN2/NG2 protein-expressing glial progenitor cells in the murine CNS: isolation, differentiation, and association with radial glia.” Glia 34 (3):213–28. doi: 10.1002/glia.1055.

Dimou, L., C. Simon, F. Kirchhoff, H. Takebayashi, and M. Gotz. 2008. “Progeny of Olig2-expressing progenitors in the gray and white matter of the adult mouse cerebral cortex.” J Neurosci 28 (41):10434–42. doi: 10.1523/JNEUROSCI.2831-08.2008.

Doetsch, F., J. M. Garcia-Verdugo, and A. Alvarez-Buylla. 1997. “Cellular composition and three-dimensional organization of the subventricular germinal zone in the adult mammalian brain.” J Neurosci 17 (13):5046–61. doi: 10.1523/JNEUROSCI.17-13-05046.1997.

Fernandez, E. M., M. D. Diaz-Ceso, and M. Vilar. 2015. “Brain expressed and X-linked (Bex) proteins are intrinsically disordered proteins (IDPs) and form new signaling hubs.” PLoS One 10 (1):e0117206. doi: 10.1371/journal.pone.0117206.

Filippov, V., G. Kronenberg, T. Pivneva, K. Reuter, B. Steiner, L. P. Wang, M. Yamaguchi, H. Kettenmann, and G. Kempermann. 2003. “Subpopulation of nestin-expressing progenitor cells in the adult murine hippocampus shows electrophysiological and morphological characteristics of astrocytes.” Mol Cell Neurosci 23 (3):373–82. doi: 10.1016/s1044-7431(03)00060-5.

Fruhbeis, C., D. Frohlich, W. P. Kuo, J. Amphornrat, S. Thilemann, A. S. Saab, F. Kirchhoff, W. Mobius, S. Goebbels, K. A. Nave, A. Schneider, M. Simons, M. Klugmann, J. Trotter, and E. M. Kramer-Albers. 2013. “Neurotransmitter-triggered transfer of exosomes mediates oligodendrocyte-neuron communication.” PLoS Biol 11 (7):e1001604. doi: 10.1371/journal.pbio.1001604.

Fu, H., J. Cai, H. Clevers, E. Fast, S. Gray, R. Greenberg, M. K. Jain, Q. Ma, M. Qiu, D. H. Rowitch, C. M. Taylor, and C. D. Stiles. 2009. “A genome-wide screen for spatially restricted expression patterns identifies transcription factors that regulate glial development.” J Neurosci 29 (36):11399–408. doi: 10.1523/JNEUROSCI.0160-09.2009.

Garcia, A. D., N. B. Doan, T. Imura, T. G. Bush, and M. V. Sofroniew. 2004. “GFAP-expressing progenitors are the principal source of constitutive neurogenesis in adult mouse forebrain.” Nat Neurosci 7 (11):1233–41. doi: 10.1038/nn1340.

Ge, W. P., X. J. Yang, Z. Zhang, H. K. Wang, W. Shen, Q. D. Deng, and S. Duan. 2006. “Long-term potentiation of neuron-glia synapses mediated by Ca2+-permeable AMPA receptors.” Science 312 (5779):1533–7. doi: 10.1126/science.1124669.

Gene Ontology, Consortium. 2021. “The Gene Ontology resource: enriching a GOld mine.” Nucleic Acids Res 49 (D1):D325–D334. doi: 10.1093/nar/gkaa1113.

Grande, A., K. Sumiyoshi, A. Lopez-Juarez, J. Howard, B. Sakthivel, B. Aronow, K. Campbell, and M. Nakafuku. 2013. “Environmental impact on direct neuronal reprogramming in vivo in the adult brain.” Nat Commun 4:2373. doi: 10.1038/ncomms3373.

Gregersen, R., T. Christensen, E. Lehrmann, N. H. Diemer, and B. Finsen. 2001. “Focal cerebral ischemia induces increased myelin basic protein and growth-associated protein-43 gene transcription in peri-infarct areas in the rat brain.” Exp Brain Res 138 (3):384–92. doi: 10.1007/s002210100715.

Guo, F., Y. Maeda, J. Ma, J. Xu, M. Horiuchi, L. Miers, F. Vaccarino, and D. Pleasure. 2010. “Pyramidal neurons are generated from oligodendroglial progenitor cells in adult piriform cortex.” J Neurosci 30 (36):12036–49. doi: 10.1523/JNEUROSCI.1360-10.2010.

Guo, F., and Y. Wang. 2023. “TCF7l2, a nuclear marker that labels premyelinating oligodendrocytes and promotes oligodendroglial lineage progression.” Glia 71 (2):143–154. doi: 10.1002/glia.24249.

Guo, Y., S. Liu, X. Zhang, L. Wang, X. Zhang, A. Hao, A. Han, and J. Yang. 2014. “Sox11 promotes endogenous neurogenesis and locomotor recovery in mice spinal cord injury.” Biochem Biophys Res Commun 446 (4):830–5. doi: 10.1016/j.bbrc.2014.02.103.

Gusel’nikova, V. V., and D. E. Korzhevskiy. 2015. “NeuN As a Neuronal Nuclear Antigen and Neuron Differentiation Marker.” Acta Naturae 7 (2):42–7.

Hackett, A. R., S. L. Yahn, K. Lyapichev, A. Dajnoki, D. H. Lee, M. Rodriguez, N. Cammer, J. Pak, S. T. Mehta, O. Bodamer, V. P. Lemmon, and J. K. Lee. 2018. “Injury type-dependent differentiation of NG2 glia into heterogeneous astrocytes.” Exp Neurol 308:72–79. doi: 10.1016/j.expneurol.2018.07.001.

Harashima, S., Y. Wang, T. Horiuchi, Y. Seino, and N. Inagaki. 2011. “Purkinje cell protein 4 positively regulates neurite outgrowth and neurotransmitter release.” J Neurosci Res 89 (10):1519–30. doi: 10.1002/jnr.22688.

Heinrich, C., M. Bergami, S. Gascon, A. Lepier, F. Vigano, L. Dimou, B. Sutor, B. Berninger, and M. Gotz. 2014. “Sox2-Mediated Conversion of NG2 Glia into Induced Neurons in the Injured Adult Cerebral Cortex.” Stem Cell Reports 3 (6):1000–1014. doi: 10.1016/j.stemcr.2014.10.007.

Hernandez, I. H., M. Villa-Gonzalez, G. Martin, M. Soto, and M. J. Perez-Alvarez. 2021. “Glial Cells as Therapeutic Approaches in Brain Ischemia-Reperfusion Injury.” Cells 10 (7). doi: 10.3390/cells10071639.

Hesp, Z. C., R. Y. Yoseph, R. Suzuki, P. Jukkola, C. Wilson, A. Nishiyama, and D. M. McTigue. 2018. “Proliferating NG2-Cell-Dependent Angiogenesis and Scar Formation Alter Axon Growth and Functional Recovery After Spinal Cord Injury in Mice.” J Neurosci 38 (6):1366–1382. doi: 10.1523/JNEUROSCI.3953-16.2017.

Hol, E. M., and M. Pekny. 2015. “Glial fibrillary acidic protein (GFAP) and the astrocyte intermediate filament system in diseases of the central nervous system.” Curr Opin Cell Biol 32:121–30. doi: 10.1016/j.ceb.2015.02.004.

Hong, X., Y. Jian, S. Ding, J. Zhou, X. Zheng, H. Zhang, B. Zhou, C. Zhuang, J. Wan, and X. Tong. 2023. “Kir4.1 channel activation in NG2 glia contributes to remyelination in ischemic stroke.” EBioMedicine 87:104406. doi: 10.1016/j.ebiom.2022.104406.

Honsa, P., H. Pivonkova, D. Dzamba, M. Filipova, and M. Anderova. 2012. “Polydendrocytes display large lineage plasticity following focal cerebral ischemia.” PLoS One 7 (5):e36816. doi: 10.1371/journal.pone.0036816.

Honsa, P., M. Valny, J. Kriska, H. Matuskova, L. Harantova, D. Kirdajova, L. Valihrach, P. Androvic, M. Kubista, and M. Anderova. 2016. “Generation of reactive astrocytes from NG2 cells is regulated by sonic hedgehog.” Glia 64 (9):1518–31. doi: 10.1002/glia.23019.

Hrckulak, D., M. Kolar, H. Strnad, and V. Korinek. 2016. “TCF/LEF Transcription Factors: An Update from the Internet Resources.” Cancers (Basel) 8 (7). doi: 10.3390/cancers8070070.

Huang, W., N. Zhao, X. Bai, K. Karram, J. Trotter, S. Goebbels, A. Scheller, and F. Kirchhoff. 2014. “Novel NG2-CreERT2 knock-in mice demonstrate heterogeneous differentiation potential of NG2 glia during development.” Glia 62 (6):896–913. doi: 10.1002/glia.22648.

Hughes, E. G., S. H. Kang, M. Fukaya, and D. E. Bergles. 2013. “Oligodendrocyte progenitors balance growth with self-repulsion to achieve homeostasis in the adult brain.” Nat Neurosci 16 (6):668–76. doi: 10.1038/nn.3390.

Jankowski, M. P., S. L. McIlwrath, X. Jing, P. K. Cornuet, K. M. Salerno, H. R. Koerber, and K. M. Albers. 2009. “Sox11 transcription factor modulates peripheral nerve regeneration in adult mice.” Brain Res 1256:43–54. doi: 10.1016/j.brainres.2008.12.032.

Jing, X., T. Wang, S. Huang, J. C. Glorioso, and K. M. Albers. 2012. “The transcription factor Sox11 promotes nerve regeneration through activation of the regeneration-associated gene Sprr1a.” Exp Neurol 233 (1):221–32. doi: 10.1016/j.expneurol.2011.10.005.

Kanazawa, Y., M. Makino, Y. Morishima, K. Yamada, T. Nabeshima, and Y. Shirasaki. 2008. “Degradation of PEP-19, a calmodulin-binding protein, by calpain is implicated in neuronal cell death induced by intracellular Ca2+ overload.” Neuroscience 154 (2):473–81. doi: 10.1016/j.neuroscience.2008.03.044.

Kang, S. H., M. Fukaya, J. K. Yang, J. D. Rothstein, and D. E. Bergles. 2010. “NG2+ CNS glial progenitors remain committed to the oligodendrocyte lineage in postnatal life and following neurodegeneration.” Neuron 68 (4):668–81. doi: 10.1016/j.neuron.2010.09.009.

Katan, M., and A. Luft. 2018. “Global Burden of Stroke.” Semin Neurol 38 (2):208–211. doi: 10.1055/s-0038-1649503.

Khazaei, M. R., H. Halfter, F. Karimzadeh, J. H. Koo, F. L. Margolis, and P. Young. 2010. “Bex1 is involved in the regeneration of axons after injury.” J Neurochem 115 (4):910–20. doi: 10.1111/j.1471-4159.2010.06960.x.

Kimura, M. T., S. Irie, S. Shoji-Hoshino, J. Mukai, D. Nadano, M. Oshimura, and T. A. Sato. 2001. “14-3-3 is involved in p75 neurotrophin receptor-mediated signal transduction.” J Biol Chem 276 (20):17291–300. doi: 10.1074/jbc.M005453200.

Kirdajova, D., and M. Anderova. 2020. “NG2 cells and their neurogenic potential.” Curr Opin Pharmacol 50:53–60. doi: 10.1016/j.coph.2019.11.005.

Kirdajova, D., L. Valihrach, M. Valny, J. Kriska, D. Krocianova, S. Benesova, P. Abaffy, D. Zucha, R. Klassen, D. Kolenicova, P. Honsa, M. Kubista, and M. Anderova. 2021. “Transient astrocyte-like NG2 glia subpopulation emerges solely following permanent brain ischemia.” Glia 69 (11):2658–2681. doi: 10.1002/glia.24064.

Kiss, T., A. Nyul-Toth, P. Balasubramanian, S. Tarantini, C. Ahire, J. DelFavero, A. Yabluchanskiy, T. Csipo, E. Farkas, G. Wiley, L. Garman, A. Csiszar, and Z. Ungvari. 2020. “Single-cell RNA sequencing identifies senescent cerebromicrovascular endothelial cells in the aged mouse brain.” Geroscience 42 (2):429–444. doi: 10.1007/s11357-020-00177-1.

Kitazono, I., T. Hamada, T. Yoshimura, M. Kirishima, S. Yokoyama, T. Akahane, and A. Tanimoto. 2020. “PCP4/PEP19 downregulates neurite outgrowth via transcriptional regulation of Ascl1 and NeuroD1 expression in human neuroblastoma M17 cells.” Lab Invest 100 (12):1551–1563. doi: 10.1038/s41374-020-0462-z.

Koo, J. H., M. Saraswati, and F. L. Margolis. 2005. “Immunolocalization of Bex protein in the mouse brain and olfactory system.” J Comp Neurol 487 (1):1–14. doi: 10.1002/cne.20486.

Koo, J. H., M. A. Smiley, R. M. Lovering, and F. L. Margolis. 2007. “Bex1 knock out mice show altered skeletal muscle regeneration.” Biochem Biophys Res Commun 363 (2):405–10. doi: 10.1016/j.bbrc.2007.08.186.

Kriska, J., L. Janeckova, D. Kirdajova, P. Honsa, T. Knotek, D. Dzamba, D. Kolenicova, O. Butenko, M. Vojtechova, M. Capek, Z. Kozmik, M. M. Taketo, V. Korinek, and M. Anderova. 2021. “Wnt/beta-Catenin Signaling Promotes Differentiation of Ischemia-Activated Adult Neural Stem/Progenitor Cells to Neuronal Precursors.” Front Neurosci 15:628983. doi: 10.3389/fnins.2021.628983.

Kuleshov, M. V., M. R. Jones, A. D. Rouillard, N. F. Fernandez, Q. Duan, Z. Wang, S. Koplev, S. L. Jenkins, K. M. Jagodnik, A. Lachmann, M. G. McDermott, C. D. Monteiro, G. W. Gundersen, and A. Ma’ayan. 2016. “Enrichr: a comprehensive gene set enrichment analysis web server 2016 update.” Nucleic Acids Res 44 (W1):W90–7. doi: 10.1093/nar/gkw377.

Lang, J., Y. Maeda, P. Bannerman, J. Xu, M. Horiuchi, D. Pleasure, and F. Guo. 2013. “Adenomatous polyposis coli regulates oligodendroglial development.” J Neurosci 33 (7):3113–30. doi: 10.1523/JNEUROSCI.3467-12.2013.

Larson, V. A., Y. Mironova, K. G. Vanderpool, A. Waisman, J. E. Rash, A. Agarwal, and D. E. Bergles. 2018. “Oligodendrocytes control potassium accumulation in white matter and seizure susceptibility.” Elife 7. doi: 10.7554/eLife.34829.

Larson, V. A., Y. Zhang, and D. E. Bergles. 2016. “Electrophysiological properties of NG2(+) cells: Matching physiological studies with gene expression profiles.” Brain Res 1638 (Pt B):138–160. doi: 10.1016/j.brainres.2015.09.010.

Li, R., P. Zhang, M. Zhang, and Z. Yao. 2020. “The roles of neuron-NG2 glia synapses in promoting oligodendrocyte development and remyelination.” Cell Tissue Res 381 (1):43–53. doi: 10.1007/s00441-020-03195-9.

Li, Y., J. Wang, Y. Zheng, Y. Zhao, M. Guo, Y. Li, Q. Bao, Y. Zhang, L. Yang, and Q. Li. 2012. “Sox11 modulates neocortical development by regulating the proliferation and neuronal differentiation of cortical intermediate precursors.” Acta Biochim Biophys Sin (Shanghai) 44 (8):660–8. doi: 10.1093/abbs/gms045.

Lillien, L. E., M. Sendtner, and M. C. Raff. 1990. “Extracellular matrix-associated molecules collaborate with ciliary neurotrophic factor to induce type-2 astrocyte development.” J Cell Biol 111 (2):635–44. doi: 10.1083/jcb.111.2.635.

Lin, S. C., and D. E. Bergles. 2004. “Synaptic signaling between GABAergic interneurons and oligodendrocyte precursor cells in the hippocampus.” Nat Neurosci 7 (1):24–32. doi: 10.1038/nn1162.

Liu, L., L. McCullough, and J. Li. 2014. “Genetic deletion of calcium/calmodulin-dependent protein kinase kinase beta (CaMKK beta) or CaMK IV exacerbates stroke outcomes in ovariectomized (OVXed) female mice.” BMC Neurosci 15:118. doi: 10.1186/s12868-014-0118-2.

Liu, Y., Q. Miao, J. Yuan, S. Han, P. Zhang, S. Li, Z. Rao, W. Zhao, Q. Ye, J. Geng, X. Zhang, and L. Cheng. 2015. “Ascl1 Converts Dorsal Midbrain Astrocytes into Functional Neurons In Vivo.” J Neurosci 35 (25):9336–55. doi: 10.1523/JNEUROSCI.3975-14.2015.

Liu, Z., Y. Li, Y. Cui, C. Roberts, M. Lu, U. Wilhelmsson, M. Pekny, and M. Chopp. 2014. “Beneficial effects of gfap/vimentin reactive astrocytes for axonal remodeling and motor behavioral recovery in mice after stroke.” Glia 62 (12):2022–33. doi: 10.1002/glia.22723.

Mabuchi, T., K. Kitagawa, T. Ohtsuki, K. Kuwabara, Y. Yagita, T. Yanagihara, M. Hori, and M. Matsumoto. 2000. “Contribution of microglia/macrophages to expansion of infarction and response of oligodendrocytes after focal cerebral ischemia in rats.” Stroke 31 (7):1735–43. doi: 10.1161/01.str.31.7.1735.

MacFarlane, S. N., and H. Sontheimer. 1997. “Electrophysiological changes that accompany reactive gliosis in vitro.” J Neurosci 17 (19):7316–29. doi: 10.1523/JNEUROSCI.17-19-07316.1997.

MacFarlane, S. N., and H. Sontheimer. 2000. “Changes in ion channel expression accompany cell cycle progression of spinal cord astrocytes.” Glia 30 (1):39–48. doi: 10.1002/(sici)1098-1136(200003)30:1<39::aid-glia5>3.0.co;2-s.

MacKay, H., C. A. Scott, J. D. Duryea, M. S. Baker, E. Laritsky, A. E. Elson, T. Garland, Jr., M. L. Fiorotto, R. Chen, Y. Li, C. Coarfa, R. B. Simerly, and R. A. Waterland. 2019. “DNA methylation in AgRP neurons regulates voluntary exercise behavior in mice.” Nat Commun 10 (1):5364. doi: 10.1038/s41467-019-13339-3.

Madisen, L., T. A. Zwingman, S. M. Sunkin, S. W. Oh, H. A. Zariwala, H. Gu, L. L. Ng, R. D. Palmiter, M. J. Hawrylycz, A. R. Jones, E. S. Lein, and H. Zeng. 2010. “A robust and high-throughput Cre reporting and characterization system for the whole mouse brain.” Nat Neurosci 13 (1):133–40. doi: 10.1038/nn.2467.

Martens, M., A. Ammar, A. Riutta, A. Waagmeester, D. N. Slenter, K. Hanspers, A. Miller R, D. Digles, E. N. Lopes, F. Ehrhart, L. J. Dupuis, L. A. Winckers, S. L. Coort, E. L. Willighagen, C. T. Evelo, A. R. Pico, and M. Kutmon. 2021. “WikiPathways: connecting communities.” Nucleic Acids Res 49 (D1):D613–D621. doi: 10.1093/nar/gkaa1024.

Matsubara, S., T. Matsuda, and K. Nakashima. 2021. “Regulation of Adult Mammalian Neural Stem Cells and Neurogenesis by Cell Extrinsic and Intrinsic Factors.” Cells 10 (5). doi: 10.3390/cells10051145.

McCullough, L. D., S. Tarabishy, L. Liu, S. Benashski, Y. Xu, T. Ribar, A. Means, and J. Li. 2013. “Inhibition of calcium/calmodulin-dependent protein kinase kinase beta and calcium/calmodulin-dependent protein kinase IV is detrimental in cerebral ischemia.” Stroke 44 (9):2559–66. doi: 10.1161/STROKEAHA.113.001030.

Mignone, J. L., V. Kukekov, A. S. Chiang, D. Steindler, and G. Enikolopov. 2004. “Neural stem and progenitor cells in nestin-GFP transgenic mice.” J Comp Neurol 469 (3):311–24. doi: 10.1002/cne.10964.

Mizrak, D., H. M. Levitin, A. C. Delgado, V. Crotet, J. Yuan, Z. Chaker, V. Silva-Vargas, P. A. Sims, and F. Doetsch. 2019. “Single-Cell Analysis of Regional Differences in Adult V-SVZ Neural Stem Cell Lineages.” Cell Rep 26 (2):394–406 e5. doi: 10.1016/j.celrep.2018.12.044.

Mouton-Liger, F., S. Thomas, R. Rattenbach, L. Magnol, V. Larigaldie, A. Ledru, Y. Herault, C. Verney, and N. Creau. 2011. “PCP4 (PEP19) overexpression induces premature neuronal differentiation associated with Ca(2+) /calmodulin-dependent kinase II-delta activation in mouse models of Down syndrome.” J Comp Neurol 519 (14):2779–802. doi: 10.1002/cne.22651.

Nishiyama, A., L. Boshans, C. M. Goncalves, J. Wegrzyn, and K. D. Patel. 2016. “Lineage, fate, and fate potential of NG2-glia.” Brain Res 1638 (Pt B):116–128. doi: 10.1016/j.brainres.2015.08.013.

Nishiyama, A., A. Chang, and B. D. Trapp. 1999. “NG2+ glial cells: a novel glial cell population in the adult brain.” J Neuropathol Exp Neurol 58 (11):1113–24. doi: 10.1097/00005072-199911000-00001.

Pivonkova, H., J. Benesova, O. Butenko, A. Chvatal, and M. Anderova. 2010. “Impact of global cerebral ischemia on K+ channel expression and membrane properties of glial cells in the rat hippocampus.” Neurochem Int 57 (7):783–94. doi: 10.1016/j.neuint.2010.08.016.

Psachoulia, K., F. Jamen, K. M. Young, and W. D. Richardson. 2009. “Cell cycle dynamics of NG2 cells in the postnatal and ageing brain.” Neuron Glia Biol 5 (3-4):57–67. doi: 10.1017/S1740925X09990354.

Putkey, J. A., M. N. Waxham, T. R. Gaertner, K. J. Brewer, M. Goldsmith, Y. Kubota, and Q. K. Kleerekoper. 2008. “Acidic/IQ motif regulator of calmodulin.” J Biol Chem 283 (3):1401–1410. doi: 10.1074/jbc.M703831200.

Rabinstein, A. A. 2017. “Treatment of Acute Ischemic Stroke.” Continuum (Minneap Minn) 23 (1, Cerebrovascular Disease):62–81. doi: 10.1212/CON.0000000000000420.

Raff, M. C., R. H. Miller, and M. Noble. 1983. “A glial progenitor cell that develops in vitro into an astrocyte or an oligodendrocyte depending on culture medium.” Nature 303 (5916):390–6. doi: 10.1038/303390a0.

Rivers, L. E., K. M. Young, M. Rizzi, F. Jamen, K. Psachoulia, A. Wade, N. Kessaris, and W. D. Richardson. 2008. “PDGFRA/NG2 glia generate myelinating oligodendrocytes and piriform projection neurons in adult mice.” Nat Neurosci 11 (12):1392–401. doi: 10.1038/nn.2220.

Robins, S. C., E. Trudel, O. Rotondi, X. Liu, T. Djogo, D. Kryzskaya, C. W. Bourque, and M. V. Kokoeva. 2013. “Evidence for NG2-glia derived, adult-born functional neurons in the hypothalamus.” PLoS One 8 (10):e78236. doi: 10.1371/journal.pone.0078236.

Robins, S. C., A. Villemain, X. Liu, T. Djogo, D. Kryzskaya, K. F. Storch, and M. V. Kokoeva. 2013. “Extensive regenerative plasticity among adult NG2-glia populations is exclusively based on self-renewal.” Glia 61 (10):1735–47. doi: 10.1002/glia.22554.

Schneider, C. A., W. S. Rasband, and K. W. Eliceiri. 2012. “NIH Image to ImageJ: 25 years of image analysis.” Nat Methods 9 (7):671–5. doi: 10.1038/nmeth.2089.

Seri, B., J. M. Garcia-Verdugo, B. S. McEwen, and A. Alvarez-Buylla. 2001. “Astrocytes give rise to new neurons in the adult mammalian hippocampus.” J Neurosci 21 (18):7153–60. doi: 10.1523/JNEUROSCI.21-18-07153.2001.

Song, F. E., J. L. Huang, S. H. Lin, S. Wang, G. F. Ma, and X. P. Tong. 2017. “Roles of NG2-glia in ischemic stroke.” CNS Neurosci Ther 23 (7):547–553. doi: 10.1111/cns.12690.

Song, F., X. Hong, J. Cao, G. Ma, Y. Han, C. Cepeda, Z. Kang, T. Xu, S. Duan, J. Wan, and X. Tong. 2018. “Kir4.1 channels in NG2-glia play a role in development, potassium signaling, and ischemia-related myelin loss.” Commun Biol 1:80. doi: 10.1038/s42003-018-0083-x.

Stoeber, K., T. D. Tlsty, L. Happerfield, G. A. Thomas, S. Romanov, L. Bobrow, E. D. Williams, and G. H. Williams. 2001. “DNA replication licensing and human cell proliferation.” J Cell Sci 114 (Pt 11):2027–41. doi: 10.1242/jcs.114.11.2027.

Struebing, F. L., J. Wang, Y. Li, R. King, O. C. Mistretta, A. W. English, and E. E. Geisert. 2017. “Differential Expression of Sox11 and Bdnf mRNA Isoforms in the Injured and Regenerating Nervous Systems.” Front Mol Neurosci 10:354. doi: 10.3389/fnmol.2017.00354.

Stuart, T., A. Butler, P. Hoffman, C. Hafemeister, E. Papalexi, W. M. Mauck, 3rd, Y. Hao, M. Stoeckius, P. Smibert, and R. Satija. 2019. “Comprehensive Integration of Single-Cell Data.” Cell 177 (7):1888–1902 e21. doi: 10.1016/j.cell.2019.05.031.

Sueda, R., I. Imayoshi, Y. Harima, and R. Kageyama. 2019. “High Hes1 expression and resultant Ascl1 suppression regulate quiescent vs. active neural stem cells in the adult mouse brain.” Genes Dev 33 (9-10):511–523. doi: 10.1101/gad.323196.118.

Sueda, R., and R. Kageyama. 2021. “Oscillatory expression of Ascl1 in oligodendrogenesis.” Gene Expr Patterns 41:119198. doi: 10.1016/j.gep.2021.119198.

Swiss, V. A., T. Nguyen, J. Dugas, A. Ibrahim, B. Barres, I. P. Androulakis, and P. Casaccia. 2011. “Identification of a gene regulatory network necessary for the initiation of oligodendrocyte differentiation.” PLoS One 6 (4):e18088. doi: 10.1371/journal.pone.0018088.

Tai, W., W. Wu, L. L. Wang, H. Ni, C. Chen, J. Yang, T. Zang, Y. Zou, X. M. Xu, and C. L. Zhang. 2021. “In vivo reprogramming of NG2 glia enables adult neurogenesis and functional recovery following spinal cord injury.” Cell Stem Cell 28 (5):923–937 e4. doi: 10.1016/j.stem.2021.02.009.

Tamura, Y., Y. Kataoka, Y. Cui, Y. Takamori, Y. Watanabe, and H. Yamada. 2007. “Multi-directional differentiation of doublecortin- and NG2-immunopositive progenitor cells in the adult rat neocortex in vivo.” Eur J Neurosci 25 (12):3489–98. doi: 10.1111/j.1460-9568.2007.05617.x.

Tan, C. L., J. L. Plotkin, M. T. Veno, M. von Schimmelmann, P. Feinberg, S. Mann, A. Handler, J. Kjems, D. J. Surmeier, D. O’Carroll, P. Greengard, and A. Schaefer. 2013. “MicroRNA-128 governs neuronal excitability and motor behavior in mice.” Science 342 (6163):1254–8. doi: 10.1126/science.1244193.

Tanner, D. C., J. D. Cherry, and M. Mayer-Proschel. 2011. “Oligodendrocyte progenitors reversibly exit the cell cycle and give rise to astrocytes in response to interferon-gamma.” J Neurosci 31 (16):6235–46. doi: 10.1523/JNEUROSCI.5905-10.2011.

Ulbricht, E., T. Pannicke, M. Hollborn, M. Raap, I. Goczalik, I. Iandiev, W. Hartig, S. Uhlmann, P. Wiedemann, A. Reichenbach, A. Bringmann, and M. Francke. 2008. “Proliferative gliosis causes mislocation and inactivation of inwardly rectifying K(+) (Kir) channels in rabbit retinal glial cells.” Exp Eye Res 86 (2):305–13. doi: 10.1016/j.exer.2007.11.002.

Uxa, S., P. Castillo-Binder, R. Kohler, K. Stangner, G. A. Muller, and K. Engeland. 2021. “Ki-67 gene expression.” Cell Death Differ 28 (12):3357–3370. doi: 10.1038/s41418-021-00823-x.

Valny, M., P. Honsa, E. Waloschkova, H. Matuskova, J. Kriska, D. Kirdajova, P. Androvic, L. Valihrach, M. Kubista, and M. Anderova. 2018. “A single-cell analysis reveals multiple roles of oligodendroglial lineage cells during post-ischemic regeneration.” Glia 66 (5):1068–1081. doi: 10.1002/glia.23301.

Vasconcelos, F. F., and D. S. Castro. 2014. “Transcriptional control of vertebrate neurogenesis by the proneural factor Ascl1.” Front Cell Neurosci 8:412. doi: 10.3389/fncel.2014.00412.

Vierbuchen, T., A. Ostermeier, Z. P. Pang, Y. Kokubu, T. C. Sudhof, and M. Wernig. 2010. “Direct conversion of fibroblasts to functional neurons by defined factors.” Nature 463 (7284):1035–41. doi: 10.1038/nature08797.

Vigano, F., S. Schneider, M. Cimino, E. Bonfanti, P. Gelosa, L. Sironi, M. P. Abbracchio, and L. Dimou. 2016. “GPR17 expressing NG2-Glia: Oligodendrocyte progenitors serving as a reserve pool after injury.” Glia 64 (2):287–99. doi: 10.1002/glia.22929.

Vilar, M., M. Murillo-Carretero, H. Mira, K. Magnusson, V. Besset, and C. F. Ibanez. 2006. “Bex1, a novel interactor of the p75 neurotrophin receptor, links neurotrophin signaling to the cell cycle.” EMBO J 25 (6):1219–30. doi: 10.1038/sj.emboj.7601017.

Virani, S. S., A. Alonso, H. J. Aparicio, E. J. Benjamin, M. S. Bittencourt, C. W. Callaway, A. P. Carson, A. M. Chamberlain, S. Cheng, F. N. Delling, M. S. V. Elkind, K. R. Evenson, J. F. Ferguson, D. K. Gupta, S. S. Khan, B. M. Kissela, K. L. Knutson, C. D. Lee, T. T. Lewis, J. Liu, M. S. Loop, P. L. Lutsey, J. Ma, J. Mackey, S. S. Martin, D. B. Matchar, M. E. Mussolino, S. D. Navaneethan, A. M. Perak, G. A. Roth, Z. Samad, G. M. Satou, E. B. Schroeder, S. H. Shah, C. M. Shay, A. Stokes, L. B. VanWagner, N. Y. Wang, C. W. Tsao, Epidemiology American Heart Association Council on, Committee Prevention Statistics, and Subcommittee Stroke Statistics. 2021. “Heart Disease and Stroke Statistics-2021 Update: A Report From the American Heart Association.” Circulation 143 (8):e254–e743. doi: 10.1161/CIR.0000000000000950.

Wang, Y., L. Lin, H. Lai, L. F. Parada, and L. Lei. 2013. “Transcription factor Sox11 is essential for both embryonic and adult neurogenesis.” Dev Dyn 242 (6):638–53. doi: 10.1002/dvdy.23962.

Wharton, S. B., K. K. Chan, J. R. Anderson, K. Stoeber, and G. H. Williams. 2001. “Replicative Mcm2 protein as a novel proliferation marker in oligodendrogliomas and its relationship to Ki67 labelling index, histological grade and prognosis.” Neuropathol Appl Neurobiol 27 (4):305–13. doi: 10.1046/j.0305-1846.2001.00333.x.

Wilhelmsson, U., A. Pozo-Rodrigalvarez, M. Kalm, Y. de Pablo, A. Widestrand, M. Pekna, and M. Pekny. 2019. “The role of GFAP and vimentin in learning and memory.” Biol Chem 400 (9):1147–1156. doi: 10.1515/hsz-2019-0199.

Wilson, H. C., N. J. Scolding, and C. S. Raine. 2006. “Co-expression of PDGF alpha receptor and NG2 by oligodendrocyte precursors in human CNS and multiple sclerosis lesions.” J Neuroimmunol 176 (1-2):162–73. doi: 10.1016/j.jneuroim.2006.04.014.

Xie, Z., A. Bailey, M. V. Kuleshov, D. J. B. Clarke, J. E. Evangelista, S. L. Jenkins, A. Lachmann, M. L. Wojciechowicz, E. Kropiwnicki, K. M. Jagodnik, M. Jeon, and A. Ma’ayan. 2021. “Gene Set Knowledge Discovery with Enrichr.” Curr Protoc 1 (3):e90. doi: 10.1002/cpz1.90.

Yang, Q. K., J. X. Xiong, and Z. X. Yao. 2013. “Neuron-NG2 cell synapses: novel functions for regulating NG2 cell proliferation and differentiation.” Biomed Res Int 2013:402843. doi: 10.1155/2013/402843.

Young, K. M., K. Psachoulia, R. B. Tripathi, S. J. Dunn, L. Cossell, D. Attwell, K. Tohyama, and W. D. Richardson. 2013. “Oligodendrocyte dynamics in the healthy adult CNS: evidence for myelin remodeling.” Neuron 77 (5):873–85. doi: 10.1016/j.neuron.2013.01.006.

Yu, Y., Y. Chen, B. Kim, H. Wang, C. Zhao, X. He, L. Liu, W. Liu, L. M. Wu, M. Mao, J. R. Chan, J. Wu, and Q. R. Lu. 2013. “Olig2 targets chromatin remodelers to enhancers to initiate oligodendrocyte differentiation.” Cell 152 (1-2):248–61. doi: 10.1016/j.cell.2012.12.006.

Zawadzka, M., L. E. Rivers, S. P. Fancy, C. Zhao, R. Tripathi, F. Jamen, K. Young, A. Goncharevich, H. Pohl, M. Rizzi, D. H. Rowitch, N. Kessaris, U. Suter, W. D. Richardson, and R. J. Franklin. 2010. “CNS-resident glial progenitor/stem cells produce Schwann cells as well as oligodendrocytes during repair of CNS demyelination.” Cell Stem Cell 6 (6):578–90. doi: 10.1016/j.stem.2010.04.002.

Zeisel, A., A. B. Munoz-Manchado, S. Codeluppi, P. Lonnerberg, G. La Manno, A. Jureus, S. Marques, H. Munguba, L. He, C. Betsholtz, C. Rolny, G. Castelo-Branco, J. Hjerling-Leffler, and S. Linnarsson. 2015. “Brain structure. Cell types in the mouse cortex and hippocampus revealed by single-cell RNA-seq.” Science 347 (6226):1138–42. doi: 10.1126/science.aaa1934.

Zhang, S., L. Yan, C. Cui, Z. Wang, J. Wu, A. Lv, M. Zhao, B. Dong, W. Zhang, X. Guan, X. Tian, and C. Hao. 2020. “Downregulation of RRM2 Attenuates Retroperitoneal Liposarcoma Progression via the Akt/mTOR/4EBP1 Pathway: Clinical, Biological, and Therapeutic Significance.” Onco Targets Ther 13:6523–6537. doi: 10.2147/OTT.S246613.

Zhang, X., Y. Lan, J. Xu, F. Quan, E. Zhao, C. Deng, T. Luo, L. Xu, G. Liao, M. Yan, Y. Ping, F. Li, A. Shi, J. Bai, T. Zhao, X. Li, and Y. Xiao. 2019. “CellMarker: a manually curated resource of cell markers in human and mouse.” Nucleic Acids Res 47 (D1):D721–D728. doi: 10.1093/nar/gky900.

Zhang, Y., K. Chen, S. A. Sloan, M. L. Bennett, A. R. Scholze, S. O’Keeffe, H. P. Phatnani, P. Guarnieri, C. Caneda, N. Ruderisch, S. Deng, S. A. Liddelow, C. Zhang, R. Daneman, T. Maniatis, B. A. Barres, and J. Q. Wu. 2014. “An RNA-sequencing transcriptome and splicing database of glia, neurons, and vascular cells of the cerebral cortex.” J Neurosci 34 (36):11929–47. doi: 10.1523/JNEUROSCI.1860-14.2014.

Zhao, C., Y. Deng, L. Liu, K. Yu, L. Zhang, H. Wang, X. He, J. Wang, C. Lu, L. N. Wu, Q. Weng, M. Mao, J. Li, J. H. van Es, M. Xin, L. Parry, S. A. Goldman, H. Clevers, and Q. R. Lu. 2016. “Dual regulatory switch through interactions of Tcf7l2/Tcf4 with stage-specific partners propels oligodendroglial maturation.” Nat Commun 7:10883. doi: 10.1038/ncomms10883.

Zhou, M., G. P. Schools, and H. K. Kimelberg. 2006. “Development of GLAST(+) astrocytes and NG2(+) glia in rat hippocampus CA1: mature astrocytes are electrophysiologically passive.” J Neurophysiol 95 (1):134–43. doi: 10.1152/jn.00570.2005.

Zhu, S., M. Chen, Y. Ying, Q. Wu, Z. Huang, W. Ni, X. Wang, H. Xu, S. Bennett, J. Xiao, and J. Xu. 2022. “Versatile subtypes of pericytes and their roles in spinal cord injury repair, bone development and repair.” Bone Res 10 (1):30. doi: 10.1038/s41413-022-00203-2.

Zhu, X., R. A. Hill, D. Dietrich, M. Komitova, R. Suzuki, and A. Nishiyama. 2011. “Age-dependent fate and lineage restriction of single NG2 cells.” Development 138 (4):745–53. doi: 10.1242/dev.047951.

Zhu, X. Q., D. E. Bergles, and A. Nishiyama. 2008. “NG2 cells generate both oligodendrocytes and gray matter astrocytes.” Development 135 (1):145–157. doi: 10.1242/dev.004895.

