## Supplementary Figures for "Astrocyte-like subpopulation of NG2 glia in the adult mouse cortex exhibits characteristics of neural progenitor cells and is capable of forming neuron-like cells after ischemic injury"

**Supplementary Information**

### Supplementary Figure and Table Legends

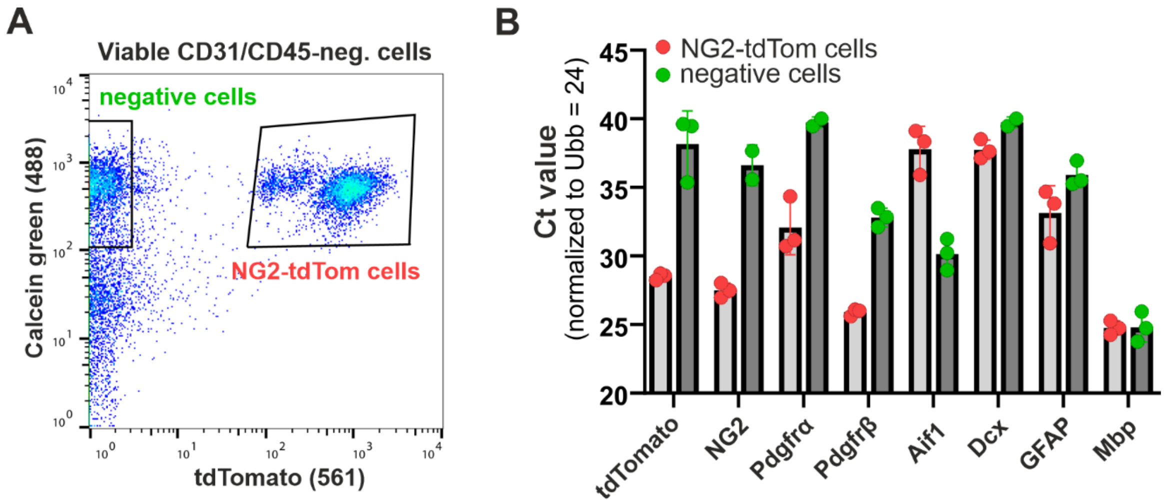


#### Supplementary Figure S1

Quantitative RT-PCR (RT-qPCR) analysis of the expression of marker genes in the cell populations sorted from sham-operated Rosa26-tdTomato/NG2/Cspg4-CreERT2 mice. (A) A representative graph shows the fluorescence-activated cell sorting (FACS) procedure to obtain viable tdTomato-positive (i.e., NG2-tdTom) and tdTomato-negative cells from the adult cortex. The viability of the cells was determined by CellTrace Calcein green fluorescence; CD31-positive endothelial cells and CD45-positive leukocytes were excluded from the sorted populations. (B) Gene expression of the selected cell type markers is shown in both NG2-tdTom and tdTomato-negative cell populations. Total RNA was isolated from three littermates. The gene expression was assessed in technical triplicates for each sample and normalized to the *Ubiquitin B* (*Ubb*) expression levels. The graph shows normalized Ct values obtained in the experiment. The individual dots represent the average value in the triplicates analyzed. The columns represent the mean of the three biological replicates. Aif1, allograft inflammatory factor 1; Dcx, doublecortin; GFAP, glial fibrillary acidic protein; Mbp, myelin basic protein; NG2, neural-glial antigen 2; Pdgfrα/β, platelet-derived growth factor receptor alpha/beta.

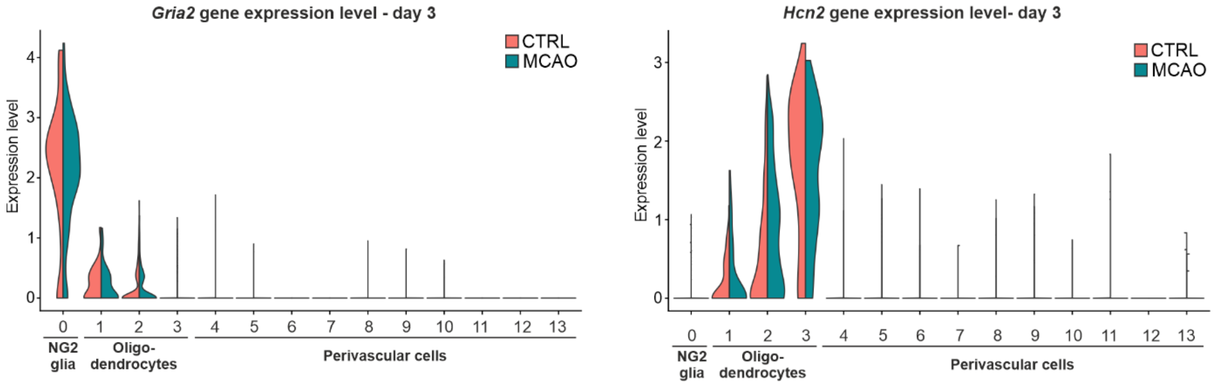


**Supplementary Figure S2**

Single-cell RNA sequencing (scRNA-seq) analysis of cells obtained from the cortex of the sham-operated (CTRL) and ischemic brain (MCAO). "Violin" plots show the expression of *Gria2* and *Hcn2* genes in the identified cell clusters 3 days upon operation. CTRL, control; Gria2, glutamate ionotropic receptor AMPA type subunit 2; Hcn2, potassium/sodium hyperpolarization-activated cyclic nucleotide-gated ion channel 2; MCAO, middle cerebral artery occlusion.

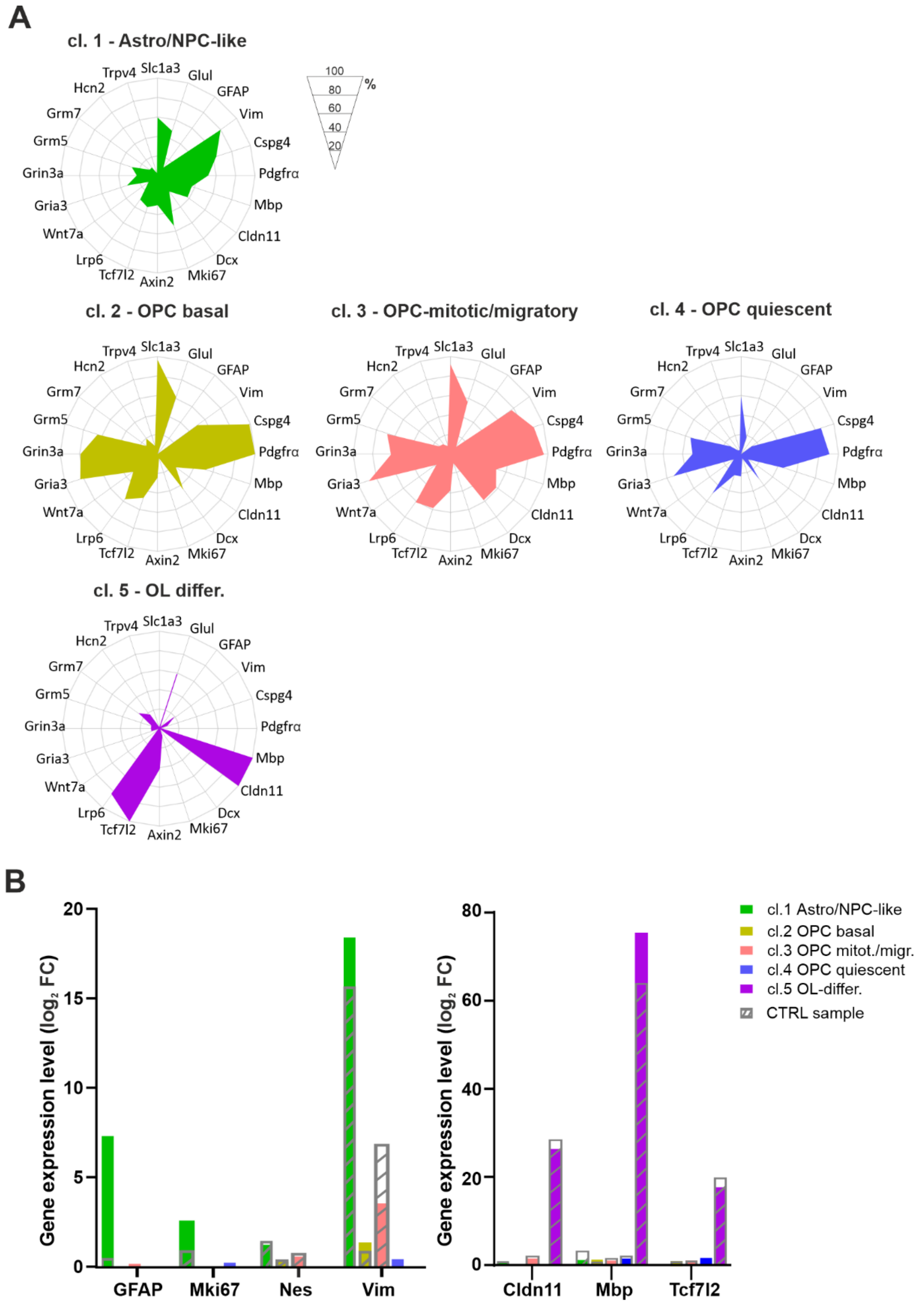


**Supplementary Figure S3**

The average expression of the specific gene set was analyzed in five NG2/Pdgfrα$^{+}$ glia subpopulations identified using the scRNA-seq analysis of the cells isolated from the healthy and ischemic cortex. NG2 glia subpopulations differ in the percentage of cells expressing the particular genes (A) and in the mRNA level of the genes (B). Gray shaded bars indicate relative gene expression in the control sample (CTRL). Cldn11, claudin 11; Cspg4, chondroitin sulfate proteoglycan 4; Dcx, doublecortin; FC, fold change; GFAP, glial fibrillary acidic protein; Glul, glutamate-ammonia ligase; Gria3, glutamate ionotropic receptor AMPA type subunit 3; Grin3a, glutamate ionotropic receptor NMDA type subunit 3A; Grm5,7, glutamate metabotropic receptor 5,7; Hcn2, potassium/sodium hyperpolarization-activated cyclic nucleotide-gated ion channel 2; Lrp6, LDL receptor related protein 6; Mbp, myelin basic protein; Mki67, marker of proliferation Ki-67; Nes, nestin; NG2, neural-glial antigen 2; NPC, neural progenitor cell; OPC, oligodendrocyte precursor cell; OL, oligodendrocyte; Pdgfrα, platelet-derived growth factor receptor alpha; Slc1a3, solute carrier family 1 member 3; Tcf7l2, transcription factor 7 like 2; Trpv4, transient receptor potential cation channel subfamily V member 4; Vim, vimentin; Wnt7a, Wnt family member 7A.

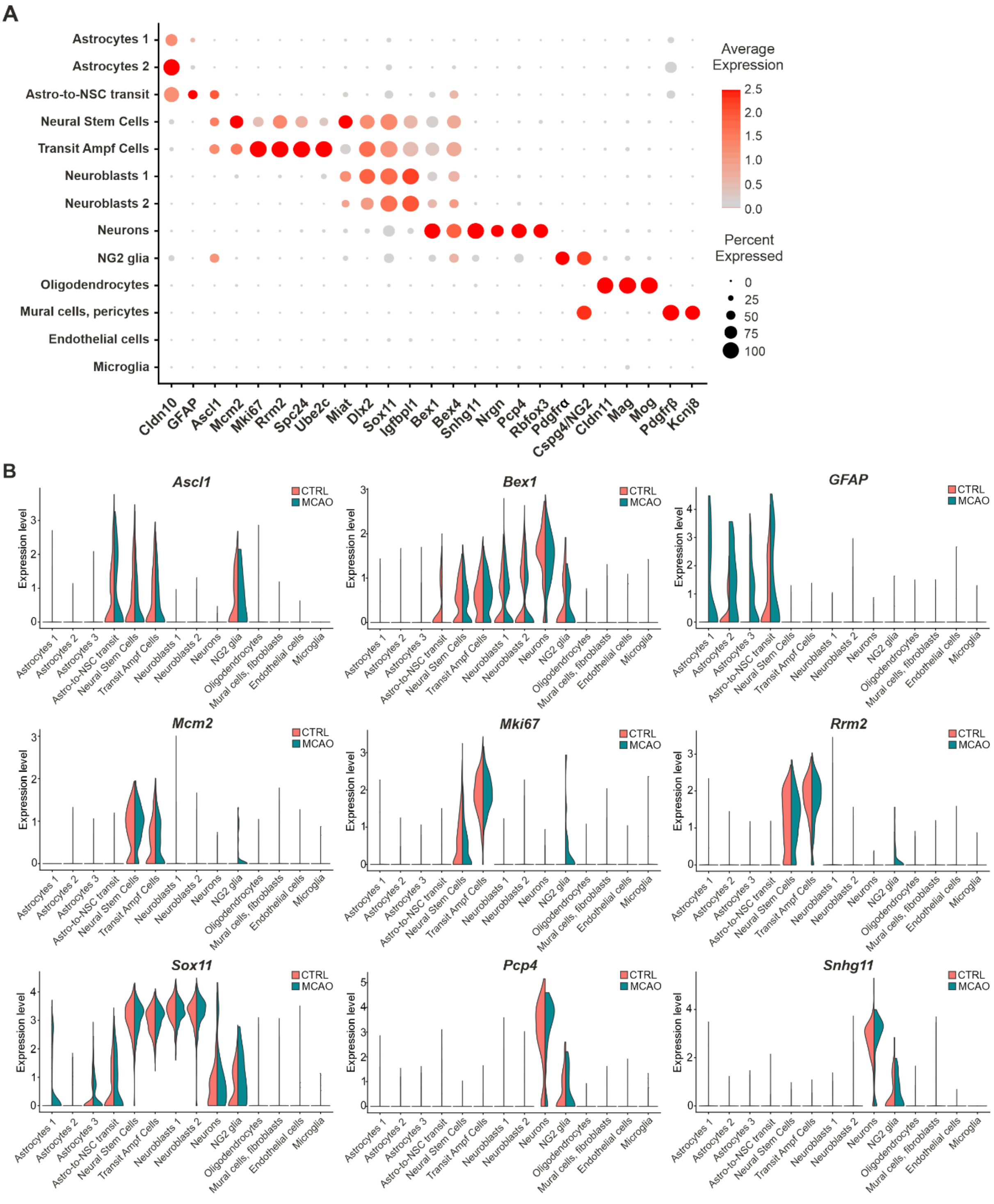


**Supplementary Figure S4**

The expression of the multiple cell type-specific genes was analyzed in the cell populations identified in the scRNA-seq from the SVZ and striatum of the sham-operated adult brain (A, dot plot) and compared with the gene expression in the corresponding region of the ischemic brain 3 days after MCAO (B, "violin" plots). Color coding reflects the increase in gene expression relative to the average gene expression across all clusters; dot size corresponds to the percentage of cells expressing the selected gene within the cluster. Ascl1, achaete-scute family BHLH transcription factor 1; Bex1,4, brain expressed X-linked 1,4; Cldn10, claudin 10; Cspg4, chondroitin sulfate proteoglycan 4; Dlx2, distal-less homeobox 2; GFAP, glial fibrillary acidic protein; Igfbpl1, insulin like growth factor binding protein like 1; Kcnj8, potassium inwardly rectifying channel subfamily J member 8; Mag, myelin Associated Glycoprotein; MCAO, middle cerebral artery occlusion; Mcm2, minichromosome maintenance complex component 2; Miat, myocardial infarction associated transcript; Mki67, marker of proliferation Ki-67; Mog, myelin oligodendrocyte glycoprotein; Nrgn, neurogranin; Pcp4, Purkinje cell protein 4; Pdgfrα,β, platelet-derived growth factor receptor alpha/beta; Rbfox3, RNA binding Fox-1 homolog 3; Rrm2, ribonucleotide reductase regulatory subunit M2; Sox11, SRY-box transcription factor 11; Snhg11, small nucleolar RNA host gene 11; Spc24, SPC24 *c*omponent of NDC80 kinetochore complex; SVZ, subventricular zone; Ube2c; ubiquitin conjugating enzyme E2 C.

**Supplementary Table S1**

Sequences of primers used in the RT-qPCR experiments.

**Supplementary Table S2**

The genes specifically expressed in each identified cluster in both CTRL and MCAO samples are listed in the corresponding sheet. Genes with the average expression $|\log_2 FC| > 0.25$ and p-value $< 0.05$ were considered as significant. Bonferroni correction based on the total number of genes in the dataset was used to calculate the p-value. The percentage of cells with the gene expression in the first and second most significant group is indicated.

**Supplementary Table S3**

The genes specifically expressed in each identified cluster of NG2 glia in both CTRL and MCAO samples at day 3 are listed in the corresponding sheet. Genes with the average expression $|\log_2 FC| > 0.25$ and p-value $< 0.05$ were considered as significant. Bonferroni correction based on the total number of genes in the dataset was used to calculate the p-value. The percentage of cells with the gene expression in the first and second most significant group is indicated.

**Supplementary Table S4**

List of genes that are expressed in the NPC-like cluster (Cl.1) of NG2 glia and simultaneously in neural stem cells (NSCs) and transit amplifying cells (TACs) identified in the subventricular zone (SVZ) and striatum of the control mice.
